# A comprehensive functional landscape of α-tubulin TUBA1A variants illuminates microtubule biology and refines clinical classification

**DOI:** 10.1101/2025.09.29.679168

**Authors:** Kaiming Xu, Zhengyang Guo, Zihan Chen, Ming Li, Zongxian Chen, Deyang Zhang, Yang Wang, Yongping Chai, Jinxiang Zhang, Hui Wang, Wei Li, Guangshuo Ou

**Affiliations:** Tsinghua-Peking Center for Life Sciences, Beijing Frontier Research Center for Biological Structure, McGovern Institute for Brain Research, State Key Laboratory of Membrane Biology, School of Life Sciences and MOE Key Laboratory for Protein Science, Tsinghua University, Beijing, China; Department of Medical Genetics, School of Basic Medicine, Tongji Medical College, Huazhong University of Science and Technology, Wuhan, China; Department of Emergency Surgery, Union Hospital, Tongji Medical College, Huazhong University of Science and Technology, Wuhan, China; School of Medicine, Tsinghua University, Beijing, China

## Abstract

Missense variant interpretation in highly conserved, paralog-rich gene families remains a critical bottleneck for precision medicine. Here, we developed an integrated experimental-computational platform to systematically assess the functional impact of all possible missense mutations in the α-tubulin TUBA1A. Combining high-throughput comprehensive mutagenesis, high-content imaging, convolutional neural network– driven phenotyping, and machine learning–guided prediction, we quantified microtubule assembly phenotypes for every coding variant. This approach outperforms conservation-based predictors and enables functional reinterpretation of disease-associated variants. Structural mapping of the TUBA1A mutational landscape reveals distinct domains critical for GTP binding, chaperone-assisted folding or protofilament interaction, illuminating diverse mechanisms of tubulin-related diseases. Integration with ACMG-AMP guidelines demonstrates this mutational landscape improves clinical variant classification in redundant gene families. This framework is broadly applicable to other structurally conserved proteins, linking variant effect prediction to mechanistic insight and clinical translation.

## Introduction

The classification of variants of uncertain significance (VUS) remains a central challenge in clinical genetics (Starita et al., 2017; Tabet et al., 2022). While advances in high-throughput sequencing have uncovered millions of coding variants (Boycott et al., 2019; Li et al., 2022), over 90% remain functionally uncharacterized, hindering diagnostic interpretation and therapeutic decision-making. Traditional genetic strategies—such as family-based linkage studies and case-control associations—lack the resolution to dissect the molecular impact of rare or private variants, particularly those resulting in subtle functional perturbations (Janes et al., 2024; Kircher et al., 2014; Sahni et al., 2013). To meet this need, deep mutational scanning (DMS) was proposed to unbiasedly evaluate functionalities of all variants, primarily through multiplexed assays of variant effect (MAVE) (Fowler et al., 2023; Fowler and Fields, 2014; Huang et al., 2025; Majithia et al., 2016; Olvera-Leon et al., 2024; Weng et al., 2024). These approaches have enabled the construction of functional maps for disease-associated genes, informing variant classification guidelines established by the American College of Medical Genetics (ACMG) and the Association for Molecular Pathology (AMP) (Richards et al., 2015).

Traditional DMS approaches rely on generating mixed libraries consisting of compound variants through well-established methods like rtITP-based mutagenesis (Haller et al., 2016), saturation genome editing (Findlay et al., 2014), saturation prime editing (Erwood et al., 2022) and CREATE (Garst et al., 2017). These platforms often rely on simple phenotypic readouts (e.g., growth assays or flow cytometry assays) (Findlay, 2021; Weile et al., 2017), which fail to capture more nuanced cellular phenotypes such as aberrant organelle morphology or spatially restricted assembly defects. These challenges are further compounded in gene families with high sequence homology, where functional redundancy can mask the effects of individual missense mutations. Alternatively, massively parallel mutagenesis enables generation of every variant as singleton. This could be accomplished by various well-established methods including SOEing (Ho et al., 1989; Hobert, 2002), PALS (Kitzman et al., 2015), one-pot mutagenesis (Wrenbeck et al., 2016), PFunkel (Firnberg and Ostermeier, 2012), SUNi (Mighell et al., 2023), massive parallel synthesis (Beltran et al., 2025) and SMuRF (Ma et al., 2024). Nonetheless, many of them rely heavily on enzyme digestion and DNA purification, rendering them time-consuming and not streamlined for parallel generation of thousands of variants.

The tubulin family exemplifies these dilemmas. Tubulins are evolutionarily conserved cytoskeletal proteins critical for many fundamental processes including cell division and intracellular transport (Janke and Magiera, 2020; McKenna et al., 2023). Pathogenic mutations in α- and β-tubulin genes cause ‘tubulinopathies’ (Bahi-Buisson et al., 2014; Cushion et al., 2023; Tischfield et al., 2011). Despite their clinical relevance, the majority of tubulin missense variants remain unclassified. Traditional DMS approaches struggle to resolve these variants: the human genome encodes nine α-tubulin isotypes sharing over 90% sequence identity, leading to functional compensation that obscures isotype-specific pathogenicity (Janke and Magiera, 2020). Computational predictors such as AlphaMissense (Cheng et al., 2023) which heavily prioritize evolutionary conservation, assign high pathogenicity scores to nearly all α-tubulin variants, probably leading to overprediction of pathogenic effects in paralogous genes (Attard et al., 2022).

To overcome these challenges, we developed an integrative experimental-computational pipeline for functional annotation of the human α-tubulin *tuba1a*. We performed saturation mutagenesis to generate all 2,683 possible coding single-nucleotide variants (SNVs) and resolved their subcellular phenotypes within several weeks. Furthermore, a machine learning model trained on imaging data extended predictions to all possible missense mutations in TUBA1A (Greener et al., 2022). This comprehensive atlas reveals structurally and functionally constrained residues required for microtubule (MT) assembly, paving the way for accurate understanding of tubulin-related disorders, accelerated construction of human SNV atlas and development of precision medicine (Ashley, 2016; Beltran et al., 2025; Fowler et al., 2023). Our study makes three key contributions: (1) it establishes a generalized framework for variant interpretation with both accuracy and speed; (2) it yields mechanistic insights into MT assembly and tubulin structure–function relationships; (3) it provides functional evidences to support variant reclassification according to ACMG-AMP guidelines, improving the clinical resolution of tubulin-related diseases.

## Results

### Generation of a comprehensive TUBA1A coding SNV library

TUBA1A encodes an essential α-tubulin isotype and harbors the highest density of disease-associated missense mutations among all tubulin isotypes (Cushion et al., 2023; Tischfield et al., 2011). To enable visualization of tubulin variants while preserving their native functionalities, we employed a previously developed strategy that inserts GFP11 into the flexible H1-S2 loop of TUBA1A (Xu et al., 2024). Fluorescence complementation with cytosolic GFP1–10 permitted visualization of mutation-associated phenotypes (Cabantous et al., 2005) (Figure S1A).

To rapidly and cost-effectively generate all 2,683 coding SNVs of TUBA1A, we optimized an overlap extension PCR (or SOEing)-based method (Figure 1A-1C) (Ho et al., 1989; Xiao et al., 2007). First, we demonstrated that linear PCR fragments— including a CMV promoter, the green fluorescence protein (EGFP) coding sequence and a downstream 3’ untranslated region (UTR)—can be effectively expressed in HeLa cells (Figure S1B). To enable high-fidelity generation of site-specific missense mutations and eliminate wild-type template contamination, we developed an automated and tail-based mutagenesis strategy (Figure 1B-C; see Method S1 for detailed protocol). A computational pipeline was implemented to automatically design primer pairs carrying missense SNVs with uniform melting temperatures (Figure 1B; Table S1; see Methods). For each mutation, two PCR fragments are generated: one spanning from the 5’ UTR to the mutation site, and the other from the mutation site to the 3’ UTR. In the initial PCR, primers are appended with short synthetic primer tails absent from the template plasmid. These tails serve as binding sites for external primers in a subsequent fusion PCR, which seamlessly joins the two fragments. Because only fragments containing these non-template tails are amplified in the final step, residual wild-type plasmid could be excluded from the reaction (Figure 1C and S1C). The entire workflow is fully automated in 96-well blocks (or plates) and can be completed within 1-2 days, depending on thermocycler availability. Consequently, this approach can be directly linked to downstream imaging and phenotypic analyses at the cellular level. For each variant, the economic expenditure is about US$3, lower than previous methodologies.

**Figure 1.**
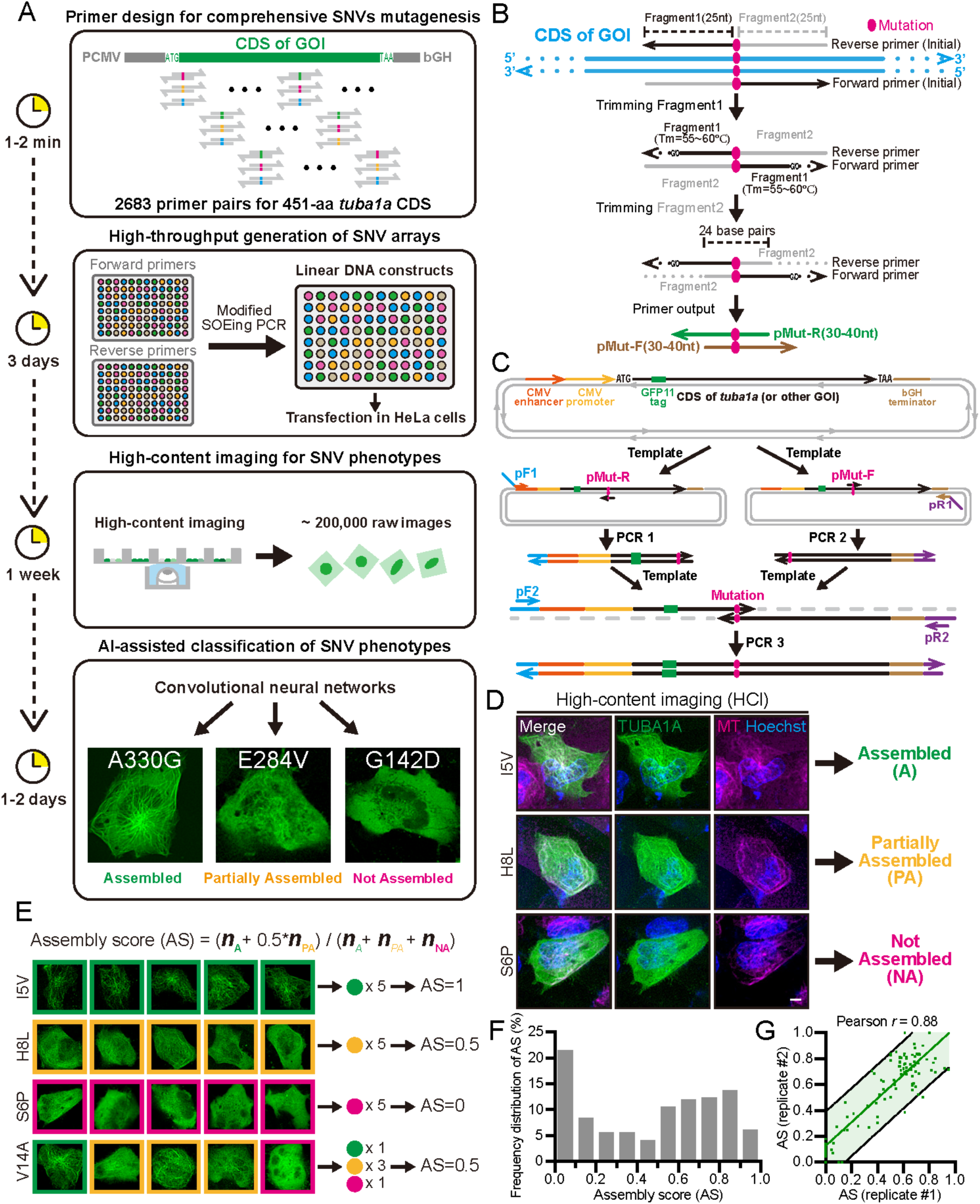
Streamlining the comprehensive phenotyping framework and calculating assembly score (AS) of TUBA1A variants. (A) Schematic overview of the comprehensive phenotyping framework. First, an automatic primer design pipeline (Figure 1B) generates variant-specific primer pairs for all coding single nucleotide variants (SNVs) of gene of interest (GOI) (time period: 1-2 min). Second, modified SOEing PCR is performed to synthesize linear DNA products in large scale (Figure 1C) (time period: 3 days). Third, high-content imaging (HCI) is performed for unbiased, automated and high-throughput acquisition of phenotypic images for each SNV. Approximately 200,000 raw images were captured for totally 2,683 TUBA1A SNVs (time period: 1-2 weeks). Finally, a deep learning-based model is trained for automatic classification of all cellular phenotypes expressing TUBA1A SNVs (Figure 1D) (time period: 1-2 days) (see Methods). (B) Workflow of automatic primer design pipeline for site-directed mutagenesis (see Methods). pMut-F and pMut-R are variant-specific primer pairs. (C) Schematic model of modified SOEing PCR (see Method S1). (D) Representative HCI images showing three distinct categories of cellular phenotypes of TUBA1A SNVs. Microtubules (MTs) are stained using 0.1x Tubulin Tracker Deep Red. A, assembled; PA, partially assembled; NA, not assembled. Scale bar, 10 μm. (E) Calculation of assembly score (AS) of each TUBA1A SNV according to its cellular phenotype (see Methods). (F) Frequency distribution of AS for all 2,683 TUBA1A SNVs. Each bar represents an interval of 0.1. (G) Correlation scatter plot of AS between two independent experimental replicates. 102 SNVs possessing variable AS values were randomly chosen.

We randomly selected 96 variants to assess the fidelity of this mutagenesis pipeline. Gel electrophoresis confirmed amplification of products at the expected sizes (Figure S1D), and Sanger sequencing covering the entire *tuba1a* CDS verified the accuracy, without undesired contamination (Figure S1E; Table S2).

### Development of an artificial intelligence (AI)-based platform for high-content imaging analysis

To systematically evaluate the functional impact of TUBA1A SNVs on MT assembly, we developed a high-content imaging (HCI) platform coupled with AI-based image analysis. HeLa cells transiently expressing split-GFP-tagged TUBA1A variants were stained with tubulin tracker before live-cell HCI (see Methods). An AI-assisted autofocus algorithm was employed to dynamically choose the optimal focal plane for imaging MT architecture (see Methods). We characterized the cellular phenotypes of all 2,683 variants within 1 week, acquiring nearly 200,000 images (Figure 1A).

To automate image analysis, a convolutional neural network (CNN)-based model was trained to perform SNV phenotype classification (Figure 1A; see Methods). Despite transient transfection, phenotype scoring remained robust as the polymerization phenotypes of tubulin variants appeared independent of their expression levels. Based on MT morphology, transfection-positive cells were classified into three phenotypic categories: assembled (A), partially assembled (PA), and not assembled (NA) (Figure 1D). Assembled (A) cells showed well-organized MT arrays co-localizing with GFP signals. NA cells exhibited diffuse and non-structured GFP fluorescence, while PA cells displayed intermediate phenotypes with partially co-localization of variants with MT networks. Cells with ambiguous classification were excluded from downstream analysis. All classifications were verified by manual review (see Methods). Variants with low transfection efficiency (< 3 cells/variant) were subjected to adaptive iterative re-screening.

Next, assembly score (AS) was defined to quantitatively evaluate the MT assembly capacity of each *tuba1a* SNV, which ranged from 0 to 1 (Figure 1E) (Caicedo et al., 2017). ‘A’ cells contributed one point, ‘PA’ cells contributed half point, and ‘NA’ cells contributed zero point. Consequently, variants composed entirely of ‘A’ cells scored 1, while those consisting solely of ‘NA’ cells scored 0. This metric enabled robust, quantitative annotation of all 2,683 TUBA1A SNVs (Figure S2; Table S3). Although quasi-continuous, the AS faithfully reflects the polymerization capacity of tubulin variants. Nearly half of the variants scored below 0.5, indicating a loss-of-function phenotype (Figure 1F). The distribution of AS values was bimodal, suggesting that many deleterious variants severely disrupt MT polymerization (Figure 1F). Moreover, a replication screen of 102 SNVs confirmed the reproducibility of AS values (Pearson’s correlation coefficient *r* = 0.88 between two replicates, Figure 1G).

To examine the credibility of our live-cell imaging-based DMS approach, four representative tubulinopathy-associated *tuba1a* SNVs were chosen for validation (Figure S3A). HeLa cells transfected with these variants were stained with tubulin tracker (0.1 μM) for 2 hours, analogous to HCI condition (Figure S3A). Through super-resolution imaging, S140G variant (AS: 0.25) partially failed to assemble into MT networks, consistent with a previous study showing its partial loss-of-function effect (Keays et al., 2007). In contrast, L286F (AS: 0.83) and R402H (AS: 0.68) variants were capable to polymerize into MTs, aligned with a previous study (Figure S3A) (Kumar et al., 2010). The localization was further confirmed by immunofluorescence staining, highlighting the fidelity of our live-cell based imaging approach (Figure S3B and S3C; see Methods). Additionally, MT plus end-binding protein EB3-mCherry was co-transfected with these variants into HeLa cells, followed by time-lapse imaging (see Methods) (Zhang et al., 2015). Astonishingly, L286F and R402H variants significantly increased MT growth rates of plus ends (Figure S3D-S3F). The two putative loss-of-function variants, V137D and S140G, had no apparent effect on EB3 velocities (Figure S3D-S3F). These results heralded a notion that while *tuba1a* SNVs of low AS cause diseases in a loss-of-function manner, those tubulinopathy-related SNVs of high AS may directly impact MT dynamics in a gain-of-function manner, which accounted for their pathogenicity (Attard et al., 2022).

Besides, we also identified a *tuba1a* variant, H393D, which disrupted MT networks in a dominant-negative manner, by manually checking HCI results for all variants (Figure S4A). This MT-depolymerizing effect of H393D variant was further confirmed by super-resolution imaging (Figure S4B and S4C). These results indicate that our live imaging-based DMS was sensitive to detect MT-depolymerizing tubulin variants.

Previously, several disease-related missense mutations in another α-tubulin isotype TUBA4A including E284G and C347Y were identified to disrupt MT networks when transfected into HeLa cells (Li et al., 2023a; Li et al., 2023b). However, these variants in TUBA1A had no apparent effect on MT networks according to HCI results. To interpret this discrepancy, E284G and C347Y variants in either TUBA1A or TUBA4A were transfected into HeLa cells at equivalent quantities (Figure S4D). Neither of the two TUBA1A variants impacted MTs, in stark contrast with TUBA4A variants (Figure S4E). Collectively, these results suggest isotype-specific disease-causing mechanisms when delving into the molecular effects of tubulin variants.

### The experimental TUBA1A landscape refines predictive models

To systematically map the functional landscape of TUBA1A, we plotted AS of all 2,683 SNVs as heatmap representation, along with residue-level average AS and variety metrics (Figure 2A; see Methods). A significant positive correlation between average AS and sequence variability (Pearson’s *r* = 0.49) supports the principle that mutations at evolutionarily conserved positions (low variability) are more likely to impair tubulin function (Beltran et al., 2025; Benegas et al., 2025). This landscape also provided an intuitive grasp of important regions along primary sequences, consistent with established structural insights (Nogales et al., 1999; Nogales et al., 1998b). These data provide a high-resolution experimental framework for interpreting mutations in functionally constrained regions of α-tubulins.

**Figure 2.**
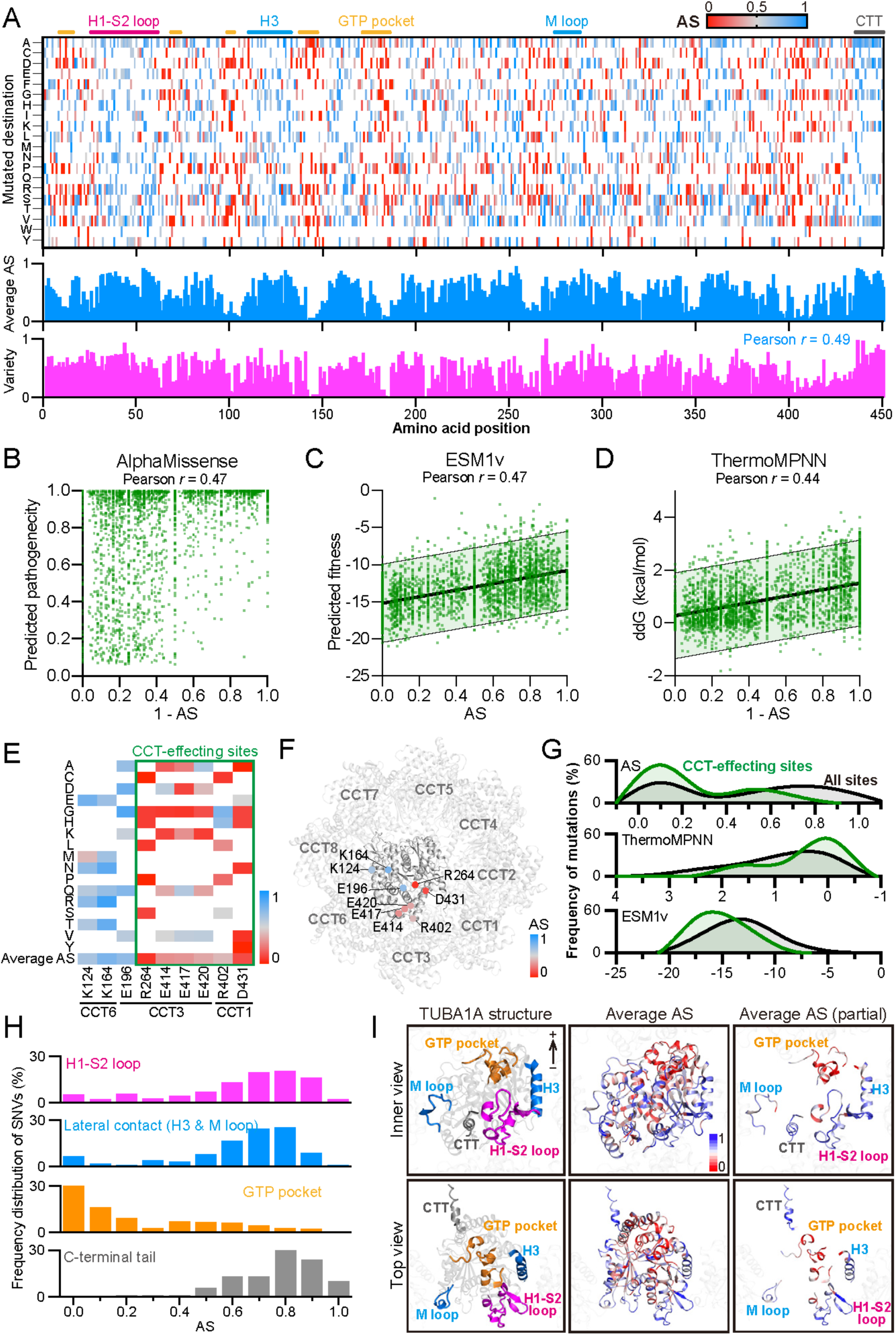
Assembly score landscape of TUBA1A highlights domain-specific functional constraints. (A) Heatmap representation of AS for all 2,683 TUBA1A coding SNVs. AS ranges from 0 (red) to 1 (blue). Well-known regions are differentially colored and indicated above the heatmap. The average AS (averaging all AS values in the same residue) and variety value (see Methods) for each residue is calculated and displayed underneath the heatmap. Pearson’s *r* = 0.49 between average AS and variety. CTT, C-terminal tail. (B) Correlation scatter plot of ‘1 - AS’ and predicted pathogenicity by AlphaMissense. (C) Correlation scatter plot of AS and predicted fitness by ESM1v. 95 % prediction band was shown. (D) Correlation scatter plot of ‘1 - AS’ and predicted ddG values by ThermoMPNN. 95 % prediction band was shown. (E) Heatmap representation of AS for CCT-interacting residues. CCT-effecting sites possess average AS below 0.5. (F) Mapping CCT-interacting residues into the TUBA1A-TRiC/CCT structural models. (G) Frequency distribution of AS (top), or predicted ddG (middle), or predicted fitness (bottom) for all 2,683 TUBA1A SNVs (gray) and CCT-effecting SNVs (green). The total percentage for each group is normalized to 100%. (H) Frequency distribution of AS for SNVs locating in H1-S2 loop (magenta), or H3/M loop (blue), or GTP-binding pocket (orange), or C-terminal tail (gray). (I) 3D structural representation of each TUBA1A functional domain and the average AS. (Left) Each domain in (H) is highlighted by different coloring. (Middle) Distribution of average AS for each residue in TUBA1A. (Right) Distribution of average AS for residues located in each functional domain. +, MT plus end.

Currently, a series of computational variant effect predictors (VEPs) have been developed to promote the functional annotation of increasing number of variants, including PolyPhen-2 (Adzhubei et al., 2010), CADD (Kircher et al., 2014), EVE (Frazer et al., 2021), ESM1v (Meier et al., 2021), ESM1b (Brandes et al., 2023), AlphaMissense (Cheng et al., 2023) and ThermoMPNN (Dieckhaus et al., 2024). To assess how well VEPs capture the functional impact of TUBA1A mutations, we compared our experimentally derived AS to predictions from three representative VEPs: AlphaMissense, ESM1v and ThermoMPNN. Among these, AlphaMissense leverages a multiple sequence alignment (MSA)–informed transformer architecture to infer pathogenicity from evolutionary constraint; ESM1v applies a large-scale protein language model to estimate functional impact based on learned sequence representations, and ThermoMPNN incorporates 3D structural information through a graph neural network to predict changes in protein thermostability.

All three predictors showed moderate correlation with experimental data (Pearson’s *r* = 0.44–0.47), supporting their overall utility. However, AlphaMissense consistently assigned high pathogenicity scores across most variants (Figure 2B), likely due to limited sequence divergence in α-tubulin MSA. Similarly, ESM1v predicted uniformly reduced fitness for a broad range of substitutions (Figure 2C; Figure S5A), reflecting strong global conservation of tubulins. In contrast, ThermoMPNN provided a more graded distribution of predicted ΔΔG° values, which did not overestimate deleterious effects (Figure 2D; Figure S5B).

These findings suggest that in highly conserved proteins, models relying on global sequence context or evolutionary conservation, such as protein language models and MSA-based predictors, may overestimate the impact of mutations due to limited sequence variability (Attard et al., 2022). In contrast, structure-informed models like ThermoMPNN, which rely on local structural context, may provide more reliable estimates of mutation effects in such cases. Critically, the experimentally derived functional landscape serves as a valuable benchmarking resource for refining and improving VEPs.

### Domain-specific functional constraints revealed by TUBA1A SNV landscape

To elucidate the structural basis of tubulin function, we analyzed how variant effects are distributed across defined functional domains of TUBA1A. Tubulin folding is mediated by a conserved chaperone-assisted pathway involving prefoldin and the TRiC/CCT complex (Gestaut et al., 2019; Gestaut et al., 2022; Vainberg et al., 1998; Yam et al., 2008). Recent cryo-EM reconstructions have pinpointed specific tubulin residues interacting with CCT subunits during the folding process (Gestaut et al., 2022). Among these residues, 6 of 9 exhibited consistently low AS upon mutation (Figure 2E–2F, termed as ‘CCT-effecting sites’), indicating their critical role in mediating chaperone engagement essential for tubulin folding. Indeed, R264C in CCT-effecting sites was associated with tubulinopathy and presumed not to fold successfully in CCT chaperones (Poirier et al., 2007; Tian et al., 2008). Moreover, ΔΔG° of SNVs in CCT-effecting sites predicted by ThermoMPNN were lower than average (Figure 2G), implying these mutations did not induce a destabilizing effect on tubulins, which further highlighted the critical impact of interaction with CCT chaperones (Dieckhaus et al., 2024). In contrast, ESM1v exhibited good performance in fitness prediction of CCT-effecting sites (Figure 2G), as it builds upon evolutionary cues involving protein-protein interaction networks (Meier et al., 2021).

We next examined AS distributions across canonical structural domains of TUBA1A (Figure 2H-2I). As expected, variants in flexible and unconserved regions—such as the H1–S2 loop and the C-terminal tail (CTT)—largely retained assembly competence, aligned with the notion that these regions are tolerant to mutations or modifications (Schatz et al., 1987; Sirajuddin et al., 2014). In contrast, mutations within the GTP-binding pocket caused severe assembly defects, underscoring the essential role of GTP/GDP dynamics in tubulin maturation and polymerization (Nogales, 2000; Rice et al., 2008). Variants in the H3 helix and M-loop—critical for lateral protofilament interactions—showed a more heterogeneous pattern: while many retained nearly normal assembly, a subset exhibited functional loss, suggesting that these domains tolerate a degree of substitution without fully disrupting tubulin polymerization.

### Plasticity of the α-tubulin GTP-binding pocket reveals context-dependent functional tolerance

GTP binding is essential for tubulin folding, dimerization, and MT dynamics (Figure 3A) (Gestaut et al., 2022; Rice et al., 2008; Roostalu et al., 2020). While β-tubulin hydrolyzes GTP at the exchangeable (E) site to regulate MT stability via the GTP cap, α-tubulin contains a non-exchangeable (N) site which binds but does not hydrolyze GTP (Alushin et al., 2014). Within the α/β-tubulin heterodimer, the GTP in N site and two coordinating loops in α-tubulin contribute directly to the intradimer face, stabilizing the overall dimer architecture (Alushin et al., 2014).

**Figure 3.**
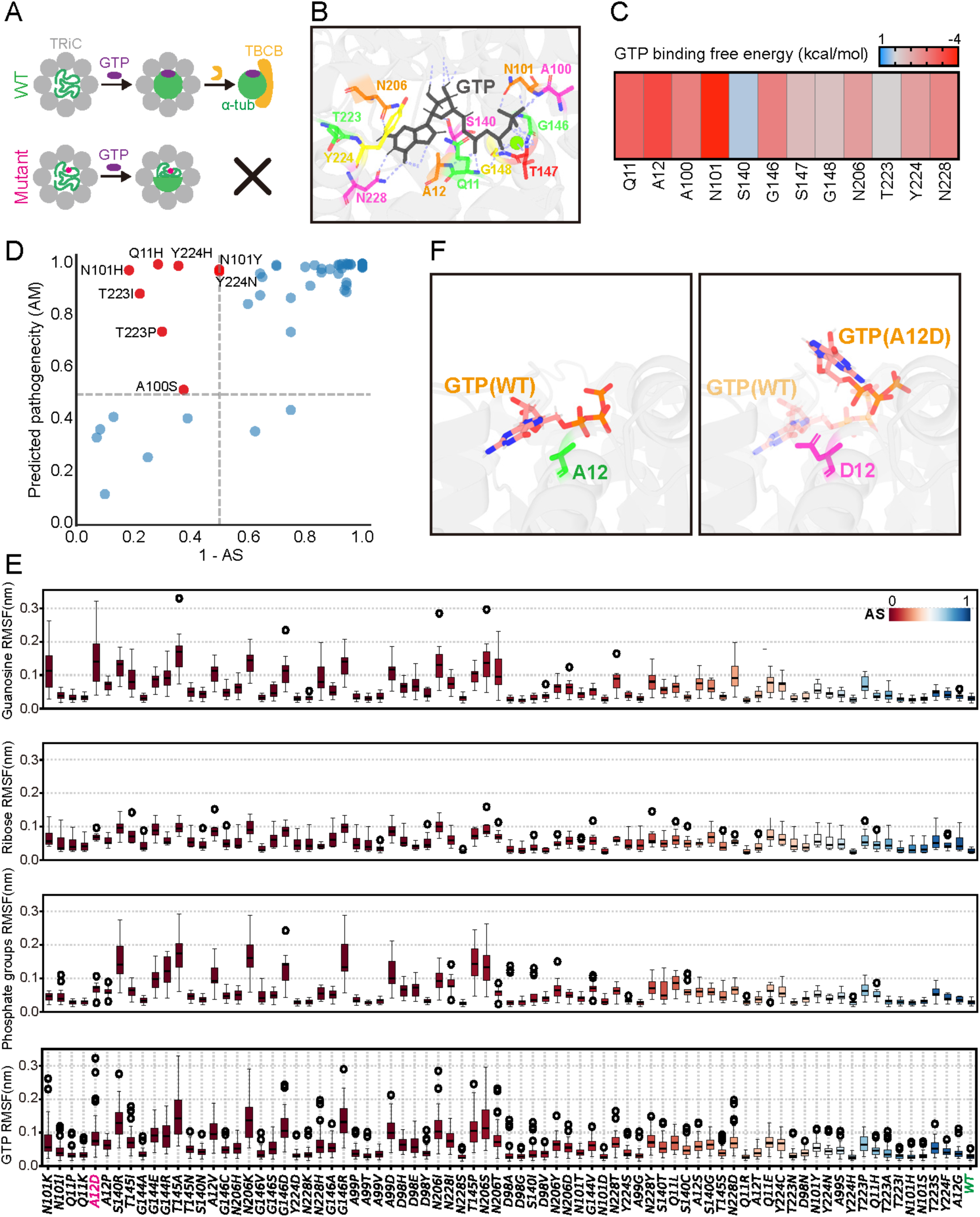
Molecular dynamics simulation (MDS) demonstrates plasticity of α-tubulin GTP-binding pocket. (A) Schematic of GTP involvement in α-tubulin maturation. Tubulin mutants (red) disrupting GTP interaction may result in compromised tubulin folding, precluding delivery of α-tubulins to downstream co-factors (TBCB). (B) Structural model of GTP-binding pocket showing critical residues in TUBA1A interacting with GTP (gray). (C) GTP binding free energy for each residue in (B) (see Methods). (D) Scatter plot of ‘1 - AS’ and predicted pathogenicity by AlphaMissense for all SNVs in (B). Variants with AS more than 0.5 and pathogenicity more than 0.5 were colored in red, otherwise blue. (E) Quantitative statistics of the root mean square fluctuation (RMSF) of distinct GTP atoms for all SNVs in (B) during MDS (see Methods). Sampling is conducted every 2 ns in the last 500 ns of the 1-μs simulation. Data are shown as median ± IQR (interquartile range). AS for each SNV is shown as gradient color ranging from red (0) to blue (1). (F) Representative status of GTP with wild-type TUBA1A (left) or A12D mutant (right) after 1-μs MDS. See also Movie S1.

To dissect the functional architecture of this GTP-binding pocket, we first identified key interacting residues based on the crystal structure of GTP-bound α-tubulin (Figure 3B and S6) (Nogales et al., 1998b). Molecular dynamics simulation (MDS) combined with MM-PBSA energy decomposition revealed that residues N101 and A12 contribute most to GTP binding free energy (Figure 3C). In contrast, residues such as Y224— despite making apparent contacts with the guanine ring in static structures—exhibited minor energetic contribution across MDS trajectories.

Comparison with experimentally derived AS revealed unexpected findings. Although AlphaMissense predicted high pathogenicity for most substitutions within this conserved pocket, several mutations—including T223I and Y224H—retained high AS (Figure 3D), consistent with their limited energetic contribution in simulations. Surprisingly, N101H also exhibited normal AS despite its prominent role in GTP coordination (Figure 3D), suggesting the compensatory side-chain interactions may preserve structural integrity. These observations underscore the limitations of conservation-based variant effect predictors in structurally complex and dynamically buffered regions.

To further probe the relationship between GTP binding stability and MT assembly, we performed MDS on 72 missense variants across 12 N-site GTP-contacting residues (Figure 3E). For each variant, the root mean square fluctuation (RMSF) of distinct GTP atoms was used to indicate the binding stability. Variants with high AS consistently showed low RMSF values, indicating stable GTP binding, while those with reduced AS such as A12D exhibited elevated RMSF, suggesting compromised GTP-tubulin coordination (Figure 3E-3F; Movie S1). These results reinforce the mechanistic link between GTP destabilization and impaired MT assembly.

Together, this work highlights the importance of integrating experimental phenotyping with atomistic simulations to resolve the complex interplay between sequence conservation, structural stability, and functional output. Besides, it demonstrates a generalizable framework for interpreting variants in ligand-binding proteins.

### Protofilament conformation and the mechanistic basis of MT assembly failure

MT plus-end dynamics are governed by the conformational properties of protofilaments, which consist of longitudinally stacked α/β-tubulin dimers (Zhang et al., 2015). Structural and computational studies have shown that protofilaments undergo coordinated twisting, bending, and rotation—modes of deformation closely coupled to MT growth, stability, and catastrophe (Fedorov et al., 2019). To mechanistically dissect how α-tubulin missense mutations impact MT assembly (Figure 4A), we developed a simulation framework to model the terminal geometry of elongating protofilaments (see Methods). This approach decomposes interdimer movements into three orthogonal components: interdimer twist (axial rotation), tangential bending and radial bending (Figure 4B) (Zhou et al., 2023).

**Figure 4.**
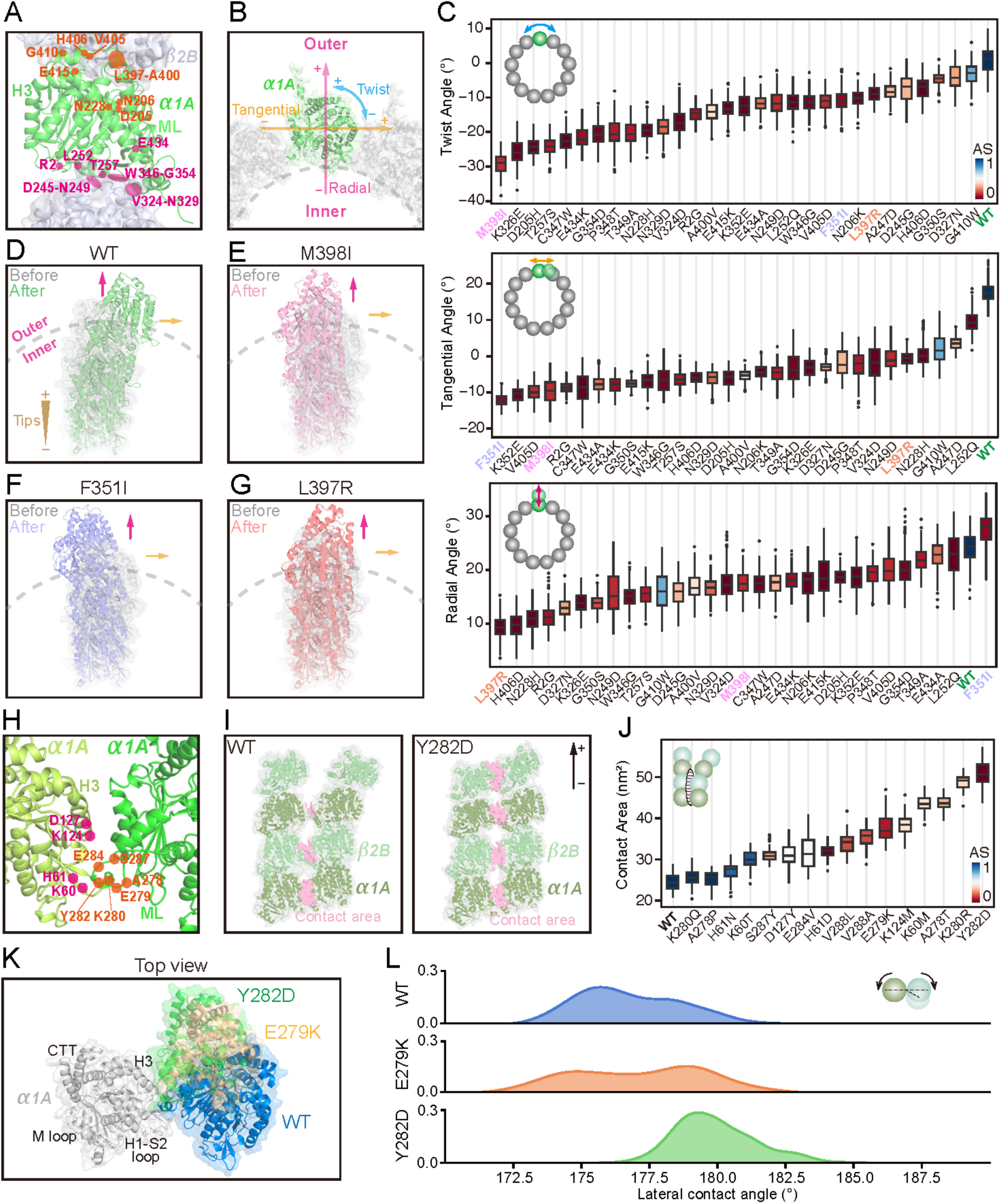
Molecular dynamics simulation demonstrates protofilament conformation affecting microtubule mechanistic properties. (A) Structural mapping of TUBA1A SNVs in intradimer or interdimer contact for MDS. TUBA1A and TUBB2B serve as the structural scaffold. H3, helix 3; ML, M loop. (B) Top view of TUBA1A (green) in MT lattice (gray). The coordinate system is defined to describe the bending angles of two neighboring tubulin heterodimers in the same PF. This includes the radial bending (magenta), tangential bending (orange) and interdimer twist (blue). (C) Quantitative statistics of twist angle (top), tangential angle (middle), or radial angle (bottom) for each SNV in (A) during MDS (see Methods). Sampling is conducted every 2 ns in the last 500 ns of the 1-μs simulation. Data are shown as median ± IQR. (D-G) Representative status of single-PF models of wild-type TUBA1A (D), M398I (E), F351I (F), or L397R (G) after MDS. Gray or colored PF represents the status before or after MDS, respectively. +, MT plus end. See also Movie S2. (H) Structural mapping of TUBA1A SNVs in lateral contact for MDS. (I) Representative status of two-PF models of TUBA1A (WT) (left) or Y282D (right) after MDS. Contact area is indicated in pink. (J) Quantitative statistics of contact area for each SNV in (H). (K) Representative status of two neighboring heterodimer models of wild-type TUBA1A, E279K or Y282D variant after simulation. Tubulin dimers on the left (gray) in each model were aligned together. (L) Frequency distribution of lateral contact angles during MDS.

Simulations of wild-type protofilaments revealed a low-energy conformational ground state characterized by minimal twist and moderate bending, consistent with cryo-EM reconstructions of growing MT ends (Figure 4C-4D) (Koning et al., 2008; McIntosh et al., 1985). By contrast, interface mutations with low AS induced marked increases in twist angle, usually accompanied by reversal of tangential bending and attenuation of radial curvature (Figure 4C-4G; Movie S2). These conformational disruptions impaired longitudinal alignment and likely resulted in tubulin disassembly. Notably, the G410W variant showed abnormal bending but retained wild-type twist and partial MT incorporation capacity, suggesting that excessive twist—more than bending—is the primary driver of assembly defect (Figure 4C).

To investigate the contribution of the α-tubulin C-terminal domain (CTD)—an evolutionary feature absent in the prokaryotic homolog FtsZ (Nogales et al., 1998a), we focused on a cluster of outward-facing residues on H11’ helix (Löwe et al., 2001), including H406, W407, and G410 (Figure S7A-S7C). Apart from most variants in H11’ helix (Figure S7B), outward-facing variants such as H406D impaired MT incorporation without disrupting monomer folding (predicted ΔΔG° = 0.5 kcal/mol, Figure S7C) or chaperone interaction, implicating altered intradimer geometry as the underlying mechanism. MDS of α/β-tubulin heterodimer revealed that H406D markedly increased root mean square deviation (RMSD) of the residue, indicating local instability of this site (Figure S7D-S7E). Trajectory analysis suggested that H406 spatially constrains W407. Disruption of this interaction, as seen in W407 mutants, similarly led to impaired MT assembly and rigid dimer intraface (Figure S7F-S7H). These findings identify helix H11’ as a previously underappreciated regulator of dimer flexibility—a functional innovation specific to eukaryotic tubulins and absent in FtsZ (Nogales et al., 1998a).

Beyond CTD, we also identified a previously less characterized region with low average AS between helix H10 and strand S9, designated Region 1 (R1) (Figure S7A). Interestingly, R1 was unique in showing poor correlation between AS and ΔΔG° (Pearson’s *r* = –0.19, Figure S7I), suggesting its functional relevance lies beyond global folding energetics. Structural mapping demonstrated R1 (residues F343–V353) localizes to the interdimer interface of the MT lattice (Figure S7J), implicating it in tubulin assembly at the MT plus end. Supporting this, two tubulinopathy-associated variants within R1 (G350V and K352M) exhibited severe MT assembly defects, with AS values of 0.18 and 0.04, respectively. These findings uncover a previously unrecognized functional hotspot, where disruption of interdimer interactions may represent a distinct pathogenic mechanism in tubulin-related disorders.

### Lateral interaction rigidity affects tubulin incorporation

Although weaker than longitudinal contacts, lateral interactions between protofilaments play a crucial role in MT stability, particularly at the growing plus-ends where lattice closure and curvature coordination are essential (Zhang et al., 2015). To evaluate how α-tubulin missense mutations affect these lateral interfaces (Figure 4H), we developed a two-protofilament MDS system in which the base α-tubulin was positionally restrained, to mimic the tip conformation of an elongating MT (see Methods). In wild-type simulations, lateral contact areas were moderate and dynamically flexible, consistent with the open and curved geometry characteristic of growing MT tips (Figure 4I) (Koning et al., 2008).

Unexpectedly, many variants with compromised MT assembly resulted in increased lateral contact area (Figure 4I-4J). To explore this apparent paradox, we employed an alternative simulation setup using two laterally adjacent tubulin dimers to analyze how alterations in lateral interface affect dimer orientation (see Methods). Strikingly, mutants such as Y282D and E279K exhibited skewed lateral contact angles centered around 180°, indicative of a rigid lateral interface (Figure 4K-4L). This contrasted with the more flexible around 176° angle observed in wild-type simulations. The loss of angular adaptability likely impairs the subtle helical pitch required for lateral closure and preventing proper lattice integration.

Together, these findings highlight that excessive lateral contact—contrary to intuition— can restrict conformational flexibility, thereby compromising the dynamic architecture necessary for tubulin incorporation. This work uncovers a previously unrecognized structural mechanism by which α-tubulin variants disrupt MT assembly—through excessive stabilization of lateral contacts that limit the flexibility required for proper tubulin integration.

### Artificial intelligence-based modeling predicts a complete tubulin functional landscape

To expand functional annotation beyond SNVs, we developed a predictive model using a graph convolutional network (GCN) trained on experimental AS (Figure 5A-5B; see Methods). This AI-driven framework generated a comprehensive *in silico* map of AS values for all possible TUBA1A missense mutations (Table S4). Model performance was validated using 52 non-SNV missense variants spanning four residues (T271, P274, P307, R390) excluded from training (Figure 5C). Predicted AS values showed strong concordance with experimental measurements (Pearson’s *r* = 0.79, Figure 6D), highlighting the accuracy and fidelity of this predictive model.

**Figure 5.**
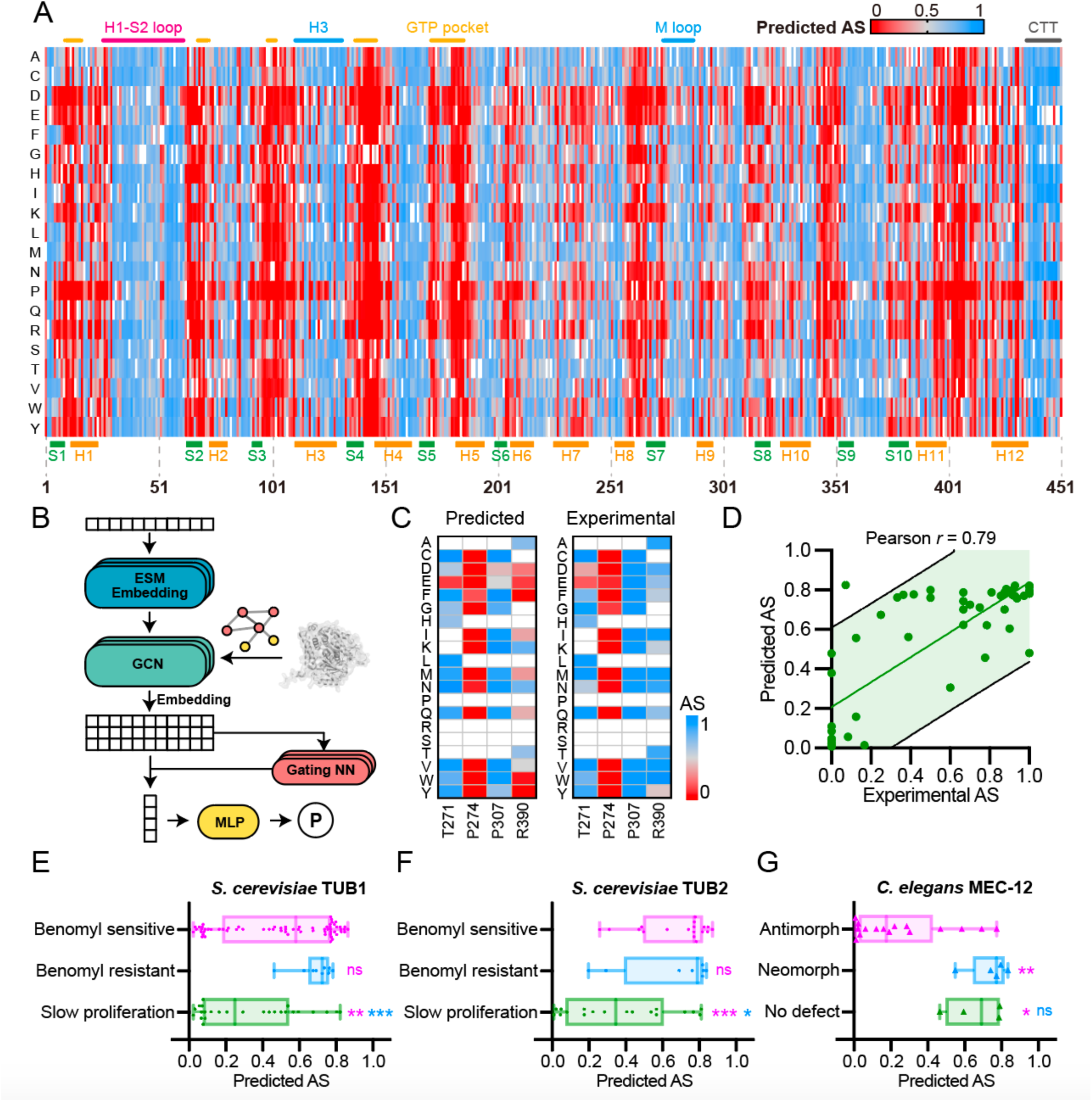
Predicting a complete functional landscape comprising all missense mutations in TUBA1A using graph convolutional network. (A) Heatmap representation showing predicted AS for all missense mutations in TUBA1A. α-helixes (H) and β-strands (S) are indicated underneath the heatmap. (B) Schematic model of AI-based AS prediction (see Methods). ESM, evolutionary scale modeling; GCN, graph convolutional network; MLP, multilayer perceptron. P, predicted AS. (C) Heatmap representation of predicted AS and experimental AS for 52 non-SNV missense mutations in TUBA1A. (D) Correlation scatter plot of experimental AS and predicted AS in (C). 95 % prediction band was shown. (E-G) Predicted AS distribution of tubulin variants in yeast (*S. cerevisiae*) TUB1 (E) or TUB2 (F), or nematode (*C. elegans)* MEC-12 (G). Data are shown as median ± IQR.

**Figure 6.**
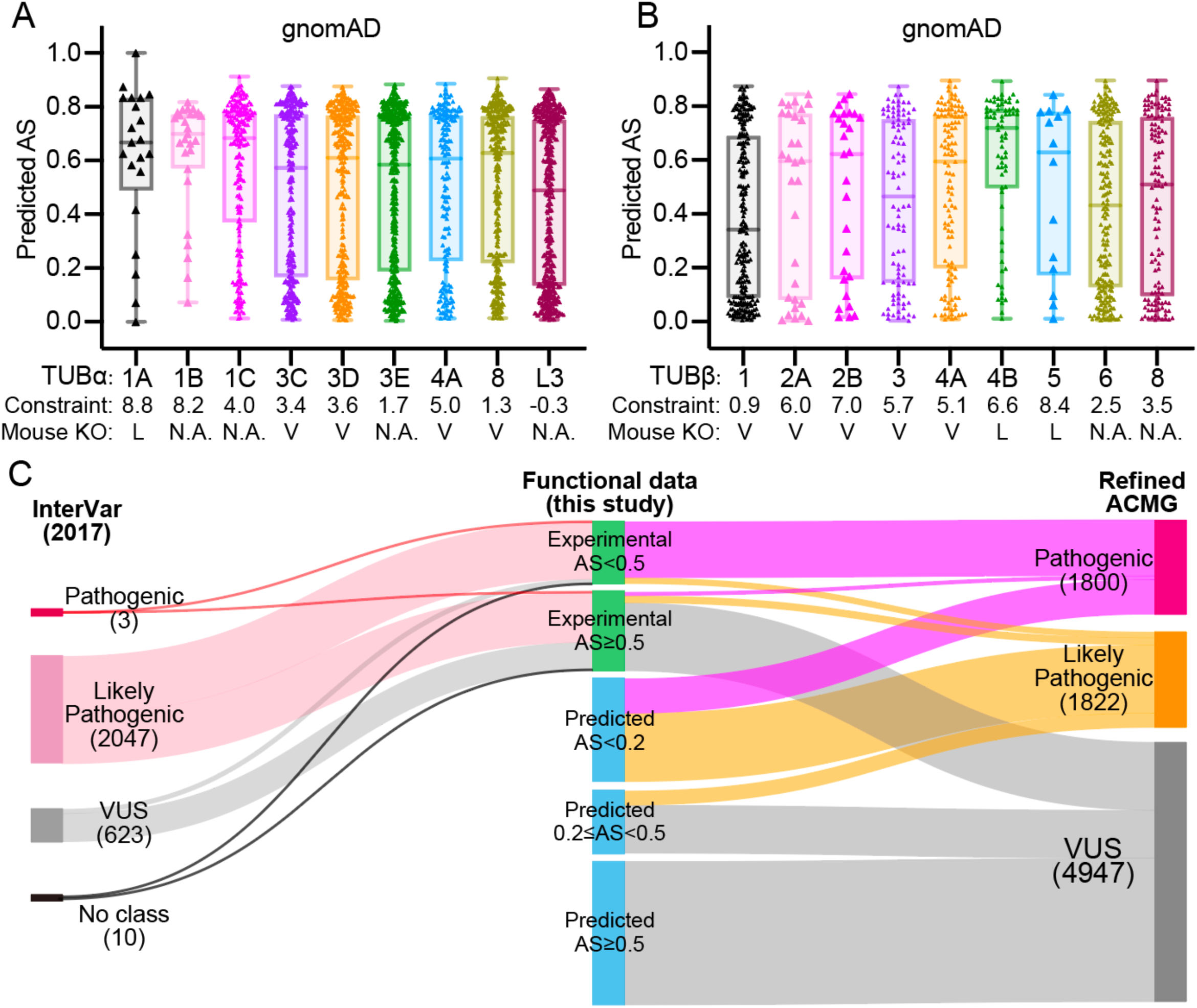
Assembly score atlas highlights differential constraints across tubulin isotypes and revises *tuba1a* clinical classification. (A and B) Predicted AS distribution of coding variants in each human α-tubulin (A) or β-tubulin (B) isotype in gnomAD database. Constraint scores are acquired from gnomAD browser (http://gnomad.broadinstitute.org/). L, lethal; V, viable; N.A., not available. Data are shown as median ± IQR. (C) Sankey plot illustrating the clinical classification of all TUBA1A coding variants before or after integration of experimental and predictive functional data into the ACMG–AMP classification framework. A previous established classification (InterVar) is presented on the left. The revised classification (Refined ACMG) is presented on the right. The numbers of variants in each classification are shown underneath the labels.

To explore the evolutionary conservation of this functional landscape, we tested whether predicted TUBA1A AS values could be extrapolated to orthologous variants in other species. In *Saccharomyces cerevisiae* where α-tubulin (TUB1/TUB3) and β-tubulin (TUB2) variants have been characterized through growth and benomyl sensitivity assays (Neff et al., 1983; Richards et al., 2000; Schatz et al., 1988; Thomas et al., 1985), homologous residues with low predicted AS were enriched among variants causing slow proliferation—consistent with loss-of-function properties (Figure 5E–5F). Conversely, most benomyl-resistant variants retained high AS, suggesting gain-of-function mutations that stabilize MTs. Differences in the AS distributions of benomyl-supersensitive mutations between TUB1 and TUB2 likely reflect the partial redundancy of α-tubulins and the essential, non-redundant role of β-tubulin in yeast (Richards et al., 2000).

In *Caenorhabditis elegans*, predicted AS values aligned with phenotypic classifications of *mec-12* variants, which encodes a mechanosensory neuron-specific α-tubulin (Lee et al., 2021; Zheng et al., 2017). Variants with antimorphic effects causing neurite growth defects exhibited significantly lower AS than neomorphic variants (associated with ectopic growth of neurites) or neutral substitutions (Figure 5G), consistent with established loss- and gain-of-function mechanisms which cause antimorph or neomorph phenotypes, respectively (Lee et al., 2021; Zheng et al., 2017).

Together, the AI-predicted TUBA1A functional landscape offers a high-resolution, scalable framework for interpreting missense variants. Its cross-species applicability underscores the evolutionary conservation of tubulins and positions this model as a generalizable tool for variant interpretation across eukaryotic systems.

### Loss-of-function landscape reveals differential constraints across tubulin isotypes

To assess how our predicted TUBA1A loss-of-function landscape reflects broader evolutionary constraint across the tubulin gene family, we assigned predicted AS to conserved missense variants in the gnomAD database (Karczewski et al., 2020), focusing on all annotated human α- and β-tubulin isotypes. Among α-tubulins, both TUBA1A (α1A) and TUBA1B (α1B) exhibited a striking depletion of low-AS variants (<0.5), consistent with their high missense constraint scores from gnomAD (α1A: 8.8; α1B: 8.2), which reflect strong selection against deleterious mutations (Figure 6A; Figure S8A–B) (Chen et al., 2024). These findings support the notion that α1A and α1B serve essential, non-redundant roles during human development.

Among β-tubulins, the most pronounced depletion of low-AS variants was observed in TUBB4B (β4B), with a modest skewing in TUBB5 (β5) (Figure 6B; Figure S8C–D). However, this pattern was not fully explained by gnomAD-based constraint scores: TUBB4B (6.6) exhibited similar levels to TUBB2A (6.0), TUBB2B (7.0), and TUBB3 (5.7)—yet these latter isotypes harbored substantially more low-AS variants. This discrepancy suggests that population-based constraint scores, which rely on mutational frequency distributions, may fail to account for certain functionally critical proteins that are captured by experimental phenotyping.

*In vivo* genetic studies corroborate these insights. Homozygous deletion of *Tubb4b* or *Tubb5* in mice results in severe postnatal lethality, underscoring their essential roles in development (Figure 6B) (Breuss et al., 2016; Dodd et al., 2024). In contrast, homozygous knockout of *Tubb2a*, *Tubb2b*, or *Tubb3* yields viable mice with no overt phenotypes (Bittermann et al., 2019; Latremoliere et al., 2018), consistent with their broader AS distributions and predicted functional tolerance. Our CRISPR-Cas9-mediated knockout experiments of *Tubb2a* and *Tubb3* similarly resulted in viable mice without apparent developmental defects (Figure S8E-S8H) (Cong et al., 2013), further validating these predictions. Together, these findings demonstrate that different tubulin isotypes vary in their functional indispensability, and experimental functional landscapes provide complementary information to population-based constraint metrics (Gao et al., 2023; Karczewski et al., 2020).

### Refining clinical classification of TUBA1A variants with functional scoring

To enhance the interpretation of TUBA1A variants in a clinical setting, we integrated functional evidences into the ACMG-AMP classification framework (Richards et al., 2015), focusing on the PS3/BS3 criteria (Brnich et al., 2019). Using experimentally derived AS, we reclassified all known TUBA1A missense SNVs and extended classification to all possible missense variants based on our predictive data (Figure 6C; Table S5; see Methods). This approach substantially increased the number of variants that could be credibly interpreted, particularly expanding the set of pathogenic and likely pathogenic classifications beyond previously established classification model such as InterVar (Li and Wang, 2017).

As part of this refinement, we reassessed the application of the PM1 criterion, which is defined for variants located in mutational hotspots or well-characterized functional domains without known benign variant. In the case of highly conserved proteins like tubulin, however, strict application of PM1 can lead to overestimation of pathogenicity due to ubiquitously evolutionarily constrained regions. To address this, we refined the application of the PM1 criterion by incorporating functional assay results: residues were assigned PM1 only if their average AS (in Figure 2A) was below 0.5, ensuring that pathogenicity assessments reflected experimentally derived functional constraints rather than conservation alone.

Importantly, variants with AS values above 0.5—reflecting normal MT assembly— were predominantly reclassified as VUS, even when located in conserved domains (Figure 6C). This phenotype-informed adjustment reduced the likelihood of false-positive pathogenic classifications and highlights the utility of AS as a quantitative, scalable metric for variant interpretation in paralog-rich, structurally conserved gene families such as tubulin.

## Discussion

The interpretation of missense variants within highly conserved and paralog-rich gene families remains a foundational challenge in clinical studies and molecular cell biology. Here, we present an integrative platform that combines comprehensive mutagenesis, high-content imaging, and AI-based modeling to generate a complete functional atlas of all coding SNVs in human α-tubulin TUBA1A.

### A scalable framework for functional annotation in conserved gene families

Fundamentally, this live-cell phenotyping platform enables subcellular-resolution profiling of MT assembly defects across thousands of variants—expanding the resolution and interpretability of multiplexed variants beyond simple phenotypes such as cell survival (Figure 1A) (Findlay, 2021; Weile et al., 2017). By incorporating CNN– based classification and an adaptive assembly score (AS), our methodology achieves both high throughput and phenotypic granularity. This framework establishes a scalable blueprint for functional annotation of essential and structurally constrained proteins.

### Mechanistic insights into tubulin structure and MT assembly

Domain-level dissection of TUBA1A variant landscape reveals strong functional constraints within GTP-binding pocket and chaperone-interaction sites. Molecular dynamics simulation links variant-induced disruption of protofilament geometry to MT assembly failure. Mutations in the α/β-intradimer face, particularly in the C-terminal domain (CTD), consistently induce compromised intradimer twist, diminished tangential and radial bending—which impede variants incorporation into MTs (Figure S7A-S7H) (Brouhard and Rice, 2018). The CTD, previously associated with microtubule-associated protein (MAP) binding and post-translational modification (Janke and Magiera, 2020), is revealed here as a key regulator of intradimer flexibility, fine-tuning the conformational adaptability of tubulins for dynamic MT assembly.

Our analysis also highlights the underappreciated role of lateral interactions in MT incorporation. Mutations such as Y282D increased lateral contact rigidity and shifted lateral contact angles toward a locked straight conformation, disrupting the helical registry required for proper lattice closure (Figure 4I-4L) (Nogales et al., 1999). These findings emphasize that both lateral and longitudinal interfaces must retain dynamic plasticity to support tubulin assembly and MT growth.

### Functional scores improve variant effect prediction and clinical classification

Comparisons with variant effect predictors demonstrate that our AS-based landscape improves conservation-based models such as AlphaMissense (Cheng et al., 2023). While sequence-based models tend to overpredict pathogenicity in conserved domains, our framework integrates subcellular phenotyping and structural modeling to distinguish variants at higher resolution. Notably, variants with high AS but predicted pathogenicity were frequently reassigned to VUS, better reflecting their benign cellular behaviors.

### Broader implications for structural biology and precision medicine

Our study refine the structural logic of tubulin function and its disease-associated perturbations. For instance, the identification of a previously uncharacterized interdimer region (R1) as a mutational hotspot expands our understanding of protofilament elongation mechanisms (Figure S7I-S7J). The convergence of low AS and clinical pathogenicity within this region illustrates how DMS uncovers novel functional domains.

Finally, the cross-species generalizability of our AI-trained model—validated within yeast and nematode orthologs—demonstrates that the rules governing tubulin function are evolutionarily conserved (Figure 5E–5G). This conservation reinforces the utility of our framework for interpreting missense mutations across both model organisms and humans, with broad relevance for developmental biology, neuroscience, and cytoskeletal research.

### Limitations of the study

This study has several limitations. While AS robustly quantifies MT polymerization capacity, it does not fully capture changes in MT dynamics. Gain-of-function effects— such as accelerated plus-end growth observed in L286F and R402H—require complementary live-cell assays to be resolved (Figure S3A-S3F). Similarly, other subtle effects may go undetected in over-expression-based systems. In vitro assays with purified tubulins may help to address these issues. Second, our screening was performed in HeLa cells, which may not fully reflect the physiological environment of TUBA1A. Isotype-specific nuances as seen in comparisons between TUBA1A and TUBA4A, underscore the importance of context-specific evaluation (Figure S4D). Furthermore, the split-GFP-based tubulin tagging strategy may impede functional analysis of H1-S2 loop, the site of GFP11 insertion. Finally, while this framework is broadly scalable, future efforts should include knock-in models to assess the impact of variants under endogenous systems. Despite these limitations, the methodology in this study offers a blueprint for functionally resolving missense variants beyond tubulins across the human genome.

### Conclusions

In conclusion, this study presents the most detailed functional atlas of a tubulin gene to date, establishing a model pipeline for systematic variant annotation in structurally conserved proteins. By bridging experimental phenotyping, structural modeling, and clinical interpretation, our platform advances both foundational and translational goals in precision medicine.

## Materials and Methods

### Plasmid construction

The cDNA construct of human *tuba1a* (NCBI sequence ID: NM_006009.4), or *tuba4a* (NM_006000.3), or *tubb8* (NM_177987.3), or *eb3* (NM_012326.4), or *gfp1-10* was cloned into pCMV3.0 vector containing upstream CMV enhancer, CMV promoter and downstream bGH terminator by homology recombination (HR) using In-Fusion® HD Cloning Kit (Takara Bio, #639650). To enable live-cell imaging of tubulin variants with high fidelity, GFP11 flanked by 16-aa GS-linkers was inserted into the G43/G44 residue pair of TUBA1A or TUBA4A, as *tuba1a (gfp11-i)* or *tuba4a (gfp11-i)* (Xu et al., 2024). The coding-domain sequence (CDS) of *tuba1a (gfp11-i)* could be found in Method S1. The DYKDDDDK (Flag) tag was inserted into the G43/G44 residue pair of TUBA1A for immunofluorescence (IF) assay. All plasmids were purified with AxyPrep Plasmid Purification Miniprep Kit (Axygen, #AP-MN-P-250).

### Automatic primer design and modified SOEing-PCR

A computational pipeline was developed to design mutagenic primers for deep mutational scanning using modified SOEing PCR (Figure 1B). The pipeline takes a target cDNA sequence, along with upstream promoter and downstream 3’ UTR regions, and systematically generates all possible coding SNVs. Tm temperature was calculated using primer3 library. The algorithm designing a single point mutation and the detailed protocol for modified SOEing-PCR are provided in Method S1.

### Transfection

HeLa cells were grown in DMEM (Gibco) containing 10 % fetal bovine serum (Yeasen) and 1 % penicillin and streptomycin (Yeasen) at 37 ℃ with 5 % CO_2_. Hela cells were seeded on 96-well glass-bottom plates (CellVis, #P96-1.5H-N) or 35 mm glass-bottom confocal dishes (CellVis, #D35C4-20-0-N) at 50-60 % confluence the day before transfection. On the second day, 150 ng of pCDNA3.0-*gfp1-10* plasmids and 1.5 μL of modified SOEing-PCR constructs containing linear *tuba1a (gfp11-i)* SNVs were co-transfected into each well of 96-well plates using polyethylenimine (PEI) MW40,000 transfection reagent (Yeasen) (Method S1). For live imaging of 35 mm confocal dishes, cells were transfected with *tuba1a (gfp11-i)* (300 ng) + *gfp1-10* (450 ng), or *tuba4a (gfp11-i)* (300 ng) + *gfp1-10* (450 ng), or EB3-mCherry (150 ng) + linear *tuba1a (gfp11-i)* SNVs (5 μL), respectively. For IF staining, flag-tagged *tuba1a* (200 ng) was tranfected into HeLa cells. The dosages of mutated constructs were consistent with their wild-type forms.

### High-content imaging (HCI)

The cells growing in 96-well glass-bottom plates were employed for HCI the day after transfection. The cells were incubated with DMEM medium containing 0.1x Tubulin Tracker^TM^ Deep Red (0.1 μM docetaxel) (Invitrogen, #T34076) at 37 °C for 30 min before live-cell imaging. Images were acquired using the ImageXpress® Confocal HT.ai system (Molecular Devices) with a 60×/1.4 objective, a 37 ℃ and 5 % CO_2_ chamber, a 60-μm pinhole camera, the autofocus mode and Cy5 / FITC channels. 36 images from different sites were acquired for each well. For each 96-well plate, HCI could be finished within 1.5-2 hours. A detailed protocol for HCI and image processing could be found in Method S1.

### Live imaging and immunofluorescence staining

The cells growing in 35 mm glass-bottom dishes were employed for imaging the day after transfection. To record EB3-mCherry movements, imaging was performed using an Axio Observer Z1 microscope (Carl Zeiss) equipped with 488 and 561 laser lines, a Yokogawa spinning disk head, an Andor iXon + EM-CCD camera, and a Zeiss 100×/1.46 objective. Images were acquired at 1 sec intervals for 1 min (EM-gain: 250, exposure time: 200 ms, 30 % of max laser) by µManager (https://www.micro-manager.org). Kymographs were generated in ImageJ by manually drawing lines along EB3-mCherry traces. For super-resolution imaging, the cells were incubated with 1x PBS (Phosphate buffered saline) containing 1x Tubulin Tracker^TM^ Deep Red and 2 μg/mL Hoechst 33342 at 37 °C for 30 min before live-cell imaging. For IF staining, the cells were fixed by 4 % paraformaldehyde (PFA) for 10 min, permeabilized by 0.1 % Triton X-100 for 10 min, and then blocked with QuickBlock™ Blocking Buffer (Beyotime) for 1 hour. Blocking buffer containing anti-DYKDDDDK 1^st^ antibody (CST, #14793) (1:800 diluted) and anti-β-tubulin 1^st^ antibody (Santa, #sc-58884) (1:200 diluted) was incubated with cells overnight at 4 ℃ to bind flag-tagged TUBA1A and microtubules (MTs) specifically. After washing with PBS for 3 times, Secondary Antibody Dilution Buffer (Beyotime) containing Dylight 488-conjugated goat-anti-rabbit 2^nd^ antibodies and Dylight 549-conjugated goat-anti-mouse 2^nd^ antibodies (Abbkine) (1:300 diluted) were incubated with cells for 1 hour at room temperature, followed by washing 3 times with PBS buffer. Imaging was performed using Zeiss LSM900 with Airyscan2 confocal microscopy (Carl Zeiss) equipped with a 63×/1.4 objective. Images were acquired at laser wavelength of 405 nm for Hoechst, 488 nm for GFP / Dylight 488, 561 nm for Dylight 549, and 640 nm for Deep Red channel. Images were taken using identical settings (Airyscan mode: Airyscan SR (super-resolution); Scan direction: bidirectional; Scan mode: frame; Scan speed: 4). Image analysis and measurement were performed with Fiji / ImageJ software (http://rsbweb.nih.gov/ij/). The visualized color of the L-561 / 640 channel was changed to colorblind-safe magenta in ImageJ. All images shown were adjusted linearly.

### Artificial intelligence-enabled phenotype classification

We employed a fine-tuned YOLO11n model for cellular phenotype classification (Redmon et al., 2016). Following HCI, images from the first plate were converted from 16-bit single-channel TIFF format (2048 × 2048 resolution) to 8-bit JPEG format (1024 × 1024 resolution) using a custom Python script based on the OpenCV library. Cellular samples were manually annotated and categorized into five distinct classes: Assembled (A), Partially Assembled (PA), Not Assembled (NA), Weak Transfection, and Dead, with approximately 100 cells labeled per class.

The model was trained on a randomly selected 80% subset of the annotated images, while the remaining 20% were used as a validation set. Training was performed over 50 epochs using the Adam optimizer with a learning rate of 0.01 and a weight decay of 0.0005.

For phenotypic analysis, images containing more than three cells not classified as ‘Weak Transfection’ or ‘Dead’ underwent further evaluation. The assembly score (AS) was subsequently quantified by a weighted average calculation based on the classification results:

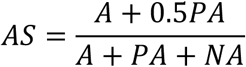

### Sequence variation estimation

We searched for sequences homologous to TUBA1A using MMseqs2 easy-search (Steinegger and Soeding, 2017), with a minimum alignment length threshold of 200 residues, querying the UniRef50 database. A total of 403 sequences were retrieved and used to evaluate the positional variation. Multiple sequence alignment (MSA) was performed using ClustalW2 (http://www.clustal.org/). The entropy at each aligned position was calculated according to the following formula:

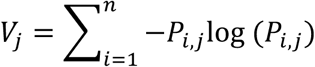

where *P_i,j_* represents the frequency of amino acid *i* at position *j* in the aligned sequences.

### Structural modeling

The amino acid sequences of TUBA1A and TUBB2B were retrieved from UniProt (http://www.uniprot.org/). Using the GMPCPP-stabilized microtubule structure (PDB ID: 3J6E) as a template, we performed homology modeling through the SWISS-MODEL workspace (Waterhouse et al., 2018). This generated three-dimensional structures for both α-tubulin (TUBA1A) and β-tubulin (TUBB2B) subunits. To simulate physiological GTP-bound conditions, we modified the GMPCPP ligand in PyMOL v2.5 (http://www.pymol.org/) by replacing the methylene group bridging the α- and β-phosphates with an oxygen atom. This conversion was achieved using PyMOL’s molecular builder toolkit with geometry optimization to maintain proper bond lengths and tetrahedral coordination angles. Ionizable residue protonation states were subsequently determined using PDBFixer (http://github.com/openmm/pdbfixer). All structural manipulations preserved the original MT lattice parameters observed in the template crystal structure.

Structural model of tubulin within TRiC/CCT complex (PDB ID: 7TUB) was downloaded from RCSB PDB database (Gestaut et al., 2022). TUBA1A was aligned to template tubulins in 7TUB using the ‘Alignment’ function of Pymol 2.5.5 for substitution. Structural model of intact MT (PDB ID: 6U42) was downloaded from RCSB PDB database (Ma et al., 2019). TUBA1A was aligned to template α-tubulins in 6U42 using the ‘Alignment’ function.

### Molecular dynamics simulation (MDS)

A virtual three-dimensional cubic reaction volume filled with TIP3P water and periodic boundary conditions was used for all simulations. The box dimensions were set such that the distance from any protein surface to the nearest box edge was at least 2 nanometers, ensuring sufficient solvation. To mimic physiological conditions, the ionic strength of the solution was adjusted to 100 mM by adding K⁺ and Cl⁻ ions, and the total charge of the system was neutralized. Simulations were performed using the GROMACS 2024.2 software package (http://www.gromacs.org/) with the CHARMM36m force field (Huang et al., 2017). Parameters for GTP were generated using the CHARMM General Force Field (CGenFF) via the CHARMM-GUI platform (http://www.charmm-gui.org/).

Each tubulin system was prepared as described in ‘Structural modeling’, followed by energy minimization using the steepest descent algorithm. Subsequently, a two-step equilibration was performed: (1) a 100 ps simulation under NVT conditions using the V-rescale thermostat (τ = 0.1 ps, T = 300 K); and (2) a 100 ps simulation under NPT conditions using the same thermostat and the C-rescale barostat (τ = 2.0 fs, P = 1 bar). During both equilibration steps, position restraints were applied to the heavy atoms of the protein and ligand. Production simulations were performed in the NPT ensemble at 300 K for 1,000 ns, using the V-rescale thermostat and C-rescale barostat. Positional restraints were applied to the Cα atoms of TUBA1A chains at the minus end of the tubulin complex throughout the production runs. Long-range electrostatic interactions were treated with the Particle Mesh Ewald method. All bonds involving hydrogen atoms were constrained using the LINCS algorithm, allowing a 2-fs integration time step. Mutations in each TUBA1A chain were introduced using the PyRosetta package with the Rosetta Energy Function 2015 (Park et al., 2016). Mutated structures were subjected to local energy minimization in PyRosetta (http://www.pyrosetta.org/) before MDS.

### Contact area calculation

The total contact area between two simulated protofilaments was estimated based on their Solvent Accessible Surface Areas (SASA). Specifically, the contact area (A) was calculated as:

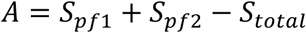

Where *S_pf_*_1_ and *S_pf_*_2_ are the SASA values of the individual protofilaments, and *S_total_* is the SASA of the protofilament complex. SASA values were computed using the GROMACS sasa module with default probe radius (0.14 nm) and parameters.

### Dimer rotation decomposition

We estimated the Euler angles describing inter-dimer rotation during a 1 μs MDS. To quantify the rotation, we selected a protofilament whose M-loop region formed a connection with an adjacent protofilament, allowing us to specifically estimate the rotation of the upper dimer relative to the lower one.

Since the tubulin dimer structure can be represented as a collection of three-dimensional atomic coordinates, the center of mass μ ∈ ℝ^1×3^ was calculated for each dimer as:

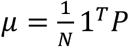

where *P* is the *N* × 3 coordinate matrix of the dimer, and 1 is an *N* × 1 vector of ones. The centered coordinates were then obtained by subtracting the center of mass:

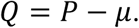

At each simulation frame *i*, the centered point cloud was defined as:

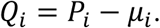

Assuming negligible internal conformational changes within each dimer, we aligned the point cloud *Q*_1_ of each frame to the initial frame *Q*_1_ through a rotation matrix *R_i_*, satisfying:

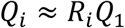

To estimate the rotation matrix *R_i_*, we used the Kabsch algorithm, which involves singular value decomposition (SVD) of the covariance matrix between *Q*_1_ and *Q*_1_:

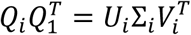

The optimal rotation matrix was then given by:

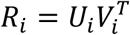

Finally, each rotation matrix *R_i_* was decomposed into Euler angles following a Z-Y-X (intrinsic) rotation sequence. The resulting angles correspond to twist (rotation about the z-axis), tangential bending (rotation about the y-axis), and radial bending (rotation about the x-axis).

### Simulation of TUBA1A-GTP-Mg^2+^ model

A TUBA1A monomer bound to GTP-Mg^2^⁺ was extracted from the constructed protofilament model described above. The system building, equilibration, and production simulation procedures followed the same protocols as for the protofilament simulations. Briefly, the CHARMM36m force field was used with TIP3P water, a 100 mM ionic strength was maintained by adding K⁺ and Cl⁻ ions, and the system was equilibrated under NVT and NPT conditions prior to production runs. For each mutant form, a 100 ns MDS was performed under the NPT ensemble at 300 K and 1 bar. After simulation, the root mean square fluctuation (RMSF) of each atom in the GTP molecule was calculated using the rmsf module in GROMACS to quantify nucleotide flexibility and assess mutation-induced changes.

### Mutation effect prediction using a graph neural network

A Graph Neural Network (GNN) was constructed to predict the mutation effects of all other 19 amino acids at each position based on our experimental data. The graph structure was derived from the AlphaFold-predicted model of TUBA1A. Specifically, Cα atom coordinates were extracted, and edges were defined between pairs of Cα atoms with a distance less than 0.8 nm.

We designed a 3-layer Graph Convolutional Network (GCN) based on this graph. Initial node embeddings were generated using ESMC and served as the input to the GCN. After graph convolution, the output node embeddings were pooled using an attention mechanism to produce a graph-level representation, which was subsequently used to predict the assembly score for each mutation (Algorithm 1).

For model training, 80 % of the experimental data were randomly selected as the training set, and the remaining 20 % were used as the validation set. The Adam optimizer was employed with a learning rate of 1×10⁻^4^ and a weight decay of 1×10^-4^. The model was trained by minimizing the mean squared error (MSE) loss, defined as *L* = (*y* – *y̅*)^2^, where *y* is the experimental score and *y̅* is the predicted score.

#### Algorithm 1

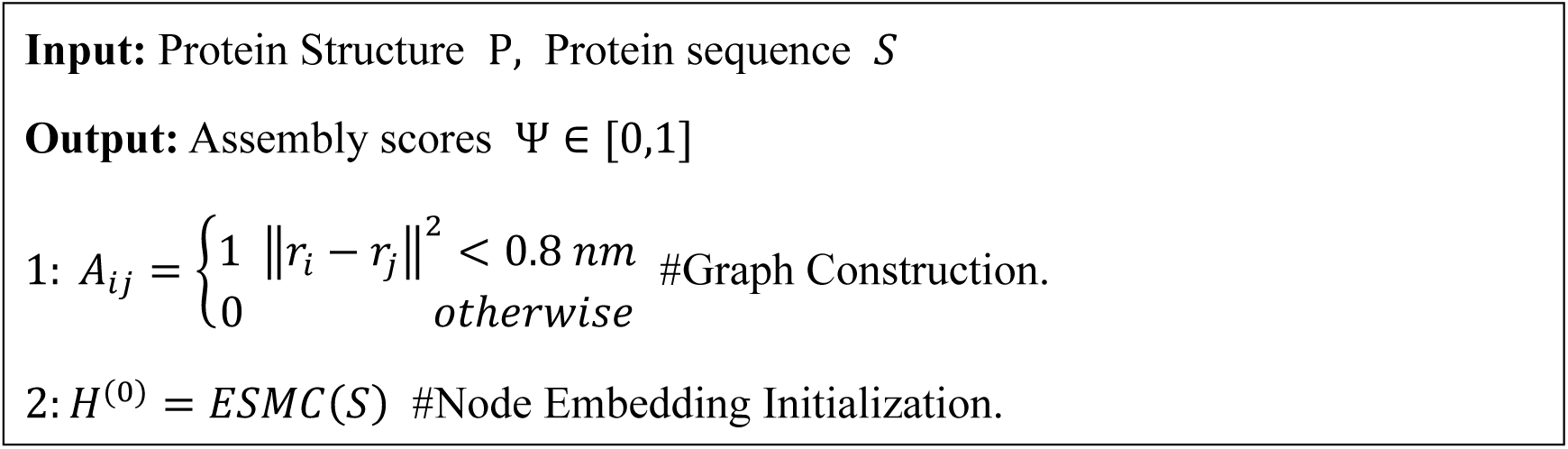
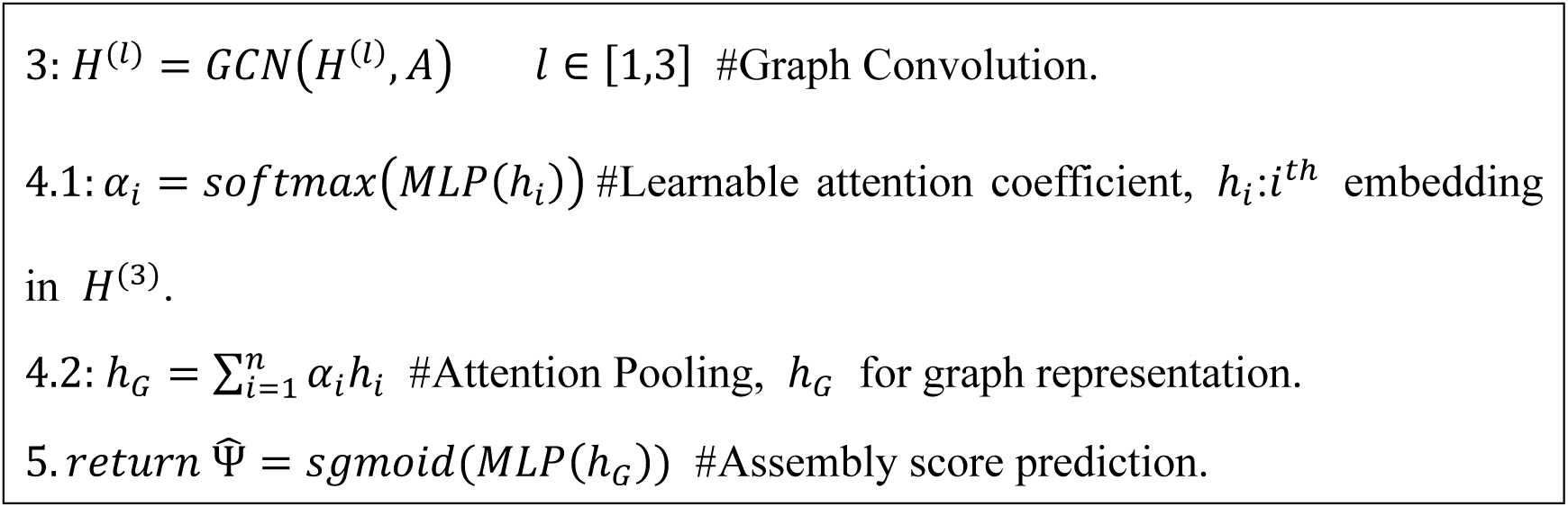

### Generation of *Tubb2a* or *Tubb3* knock-out (KO) mouse lines

All procedures for the care and use of laboratory animals was approved by the Institutional Animal Care and Use Committee (IACUC) and Internal Review Board of Tsinghua University (Animal Protocol number: 24-OGS1). *Tubb2a* (+/-) or *Tubb3* (+/-) mice were generated from wild-type C57BL/6 animals using CRISPR-Cas9 by pronuclear injection. The microinjection mix consisted of 20 ng/μL guide RNA and 20 ng/μL Cas9 mRNA. Founders were genotyped by Sanger sequencing of genomic DNA extracted from mousetail tissues, using *Tubb2a* or *Tubb3*-specific primers. Founder mice (+/-) were then crossed with C57BL/6 mice, and the heterozygous offsprings were intercrossed to produce homozygous KO lines, which were confirmed by Sanger sequencing. Consequently, frameshift mutations resulted in premature termination codons (PTCs) in *Tubb2a* or *Tubb3*, respectively. Animals were maintained in SPF environment. The guide DNA sequences used for microinjection are listed below:

For *Tubb2a* KO: GGTGGTCACTCCACTCATGG

For *Tubb3* KO: GCATGAAGAAGTGGAGACGT

### Revise the clinical classification following the ACMG-AMP guideline

Revised *tuba1a* variant classification was performed according to the standards and guidelines of ACMG-AMP (Richards et al., 2015). For established pathogenic mutations which cause human diseases, the variants were assigned PS1. Variants with experimental AS less than 0.5 or predicted AS less than 0.2 were assigned PS3, as the predicted AS was highly correlated with experimental validation. Variants at an amino acid residue where a different pathogenic missense change has been seen before, were assigned PM5. Variants at an amino acid residue whose average AS (Figure 2A) are less than 0.5 were assigned PM1. Variants with predicted AS less than 0.5 were assigned PP3. All variants in *tuba1a* were assigned PP2. Variants which are designated as ‘pathogenic’ or ‘likely pathogenic’ in ClinVar database were assigned PP5. As a result, we revised the clinical classification of all *tuba1a* coding variants (including SNVs and non SNVs) according to the rules for combining criteria (Richards et al., 2015). Notably, we revised the judging criteria for PM1 when compared to a previously established classification -- InterVar (http://wintervar.wglab.org/) (Li and Wang, 2017). According to InterVar, nearly all *tuba1a* SNVs were assigned PM1, which may hinder the distinct effects of mutational hotspots or functional domains. In our classification, residues were assigned PM1 only if their average AS was below 0.5, ensuring that pathogenicity assessments reflected experimentally derived functional characteristics.

### Quantification and statistical analysis

The sample sizes in our experiments were determined from the related published analyses. We used GraphPad Prism 9 (GraphPad Software, Inc.) for statistical analyses and calculation of Pearson’s correlation coefficient (*r*). Mann-Whitney tests (nonparametric tests) were performed to compare the differences of assembly scores between two groups. *N* represents the number of samples in each group. Statistical significances were designated as: ns *P* > 0.05, * *P* < 0.05, ** *P* < 0.01, *** *P* < 0.001 and **** *P* < 0.0001.

## Acknowledgements

This work was supported by the National Natural Science Foundation of China (NSFC) Grants 92254306 (G.O.), 323B200173 (K.X.), 32430026 (G.O.), 32021002 (G.O.), 31991191 (G.O.), and 32270721 (Y.C.). We are deeply grateful to Carsten Janke, Eva Nogales, Joseph Marsh, Hongwei Wang, Elizabeth Engle and Luke Rice for their insightful discussions and constructive feedback. We thank the Facility of Cell Imaging in the Tsinghua University Technology Center for Protein Research (especially Yuke Feng), and the Tsinghua University Laboratory Animal Resources Center (THU-LARC) (especially Yankun Yang), for their technical assistance.

## Author Contributions

G.O., K.X., Z.G. and Z.H.C. conceived and designed this study. G.O. supervised this study. K.X. and Z.H.C. performed the experiments for the establishment and implementation of the comprehensive mutagenesis framework, and the construction of TUBA1A coding SNV atlas. Z.G. performed computational analyses (deep learning-based model training, development of primer design pipeline, and molecular dynamics simulation). M.L., Z.X.C., D.Y.Z., and Y.W. assisted in computational analyses. K.X. performed data analysis and refined the clinical classification. K.X., G.O., Z.G. wrote this manuscript with the help of other authors.

## Data and Code Availability

All data and original code in this study are available upon reasonable request. All original images used in this study are publicly available.

---

**Figure S1.**
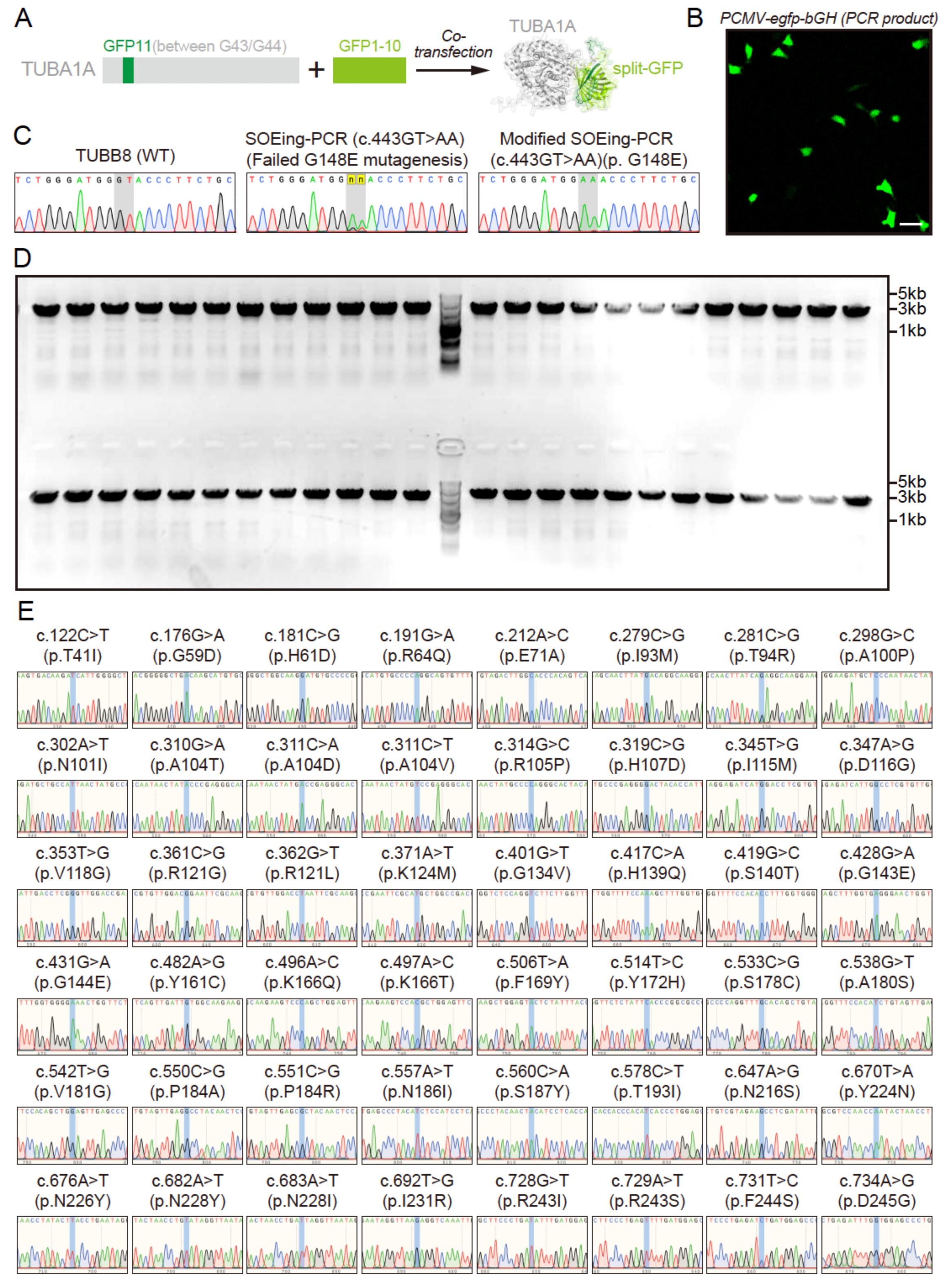
Validation of the modified SOEing-PCR, relates to Figure 1. (A) Schematic of live-cell labeling of α-tubulin TUBA1A using split-GFP technology (Xu et al., 2024). 16-aa GFP11 flanked by linkers is inserted into the G43/G44 residue pair of TUBA1A, which is co-transfected with GFP1-10 into mammalian cells. (B) Representative image of HeLa cells showing successful transfection and expression of linear *PCMV-egfp-bGH* fragments amplified by modified SOEing-PCR. Scale bar, 50 μm. (C) Peak diagrams showing Sanger sequencing results of PCR products of wild-type TUBB8 (left), or failed G148E mutagenesis by traditional SOEing-PCR (middle), or successful G148E mutagenesis by modified SOEing-PCR (right). Source nucleotides ‘GT’ (marked in gray columns) are subjected to be mutated to ‘AA’, whereas ‘GGT’ is the codon of G148 in TUBB8. (D) Agarose gel electrophoresis results of 48 representative modified SOEing-PCR products, showing successful generation of TUBA1A variants without undesired contamination. The targeted length of DNA fragment is about 3 kilobases (kb). (E) Peak diagrams showing Sanger sequencing results of PCR products of 48 representative TUBA1A variants. Source nucleotides, their positions in cDNA sequence of *tuba1a* (Method S1), and targeted mutations are indicated above each diagram (c., cDNA). Source amino acids, their positions in TUBA1A protein sequence, and targeted mutations are indicated above each diagram (p., protein). Targeted sites for mutagenesis were marked in blue columns. Sanger sequencing results of all 96 randomly selected variants could be found in Table S2.

**Figure S2.**
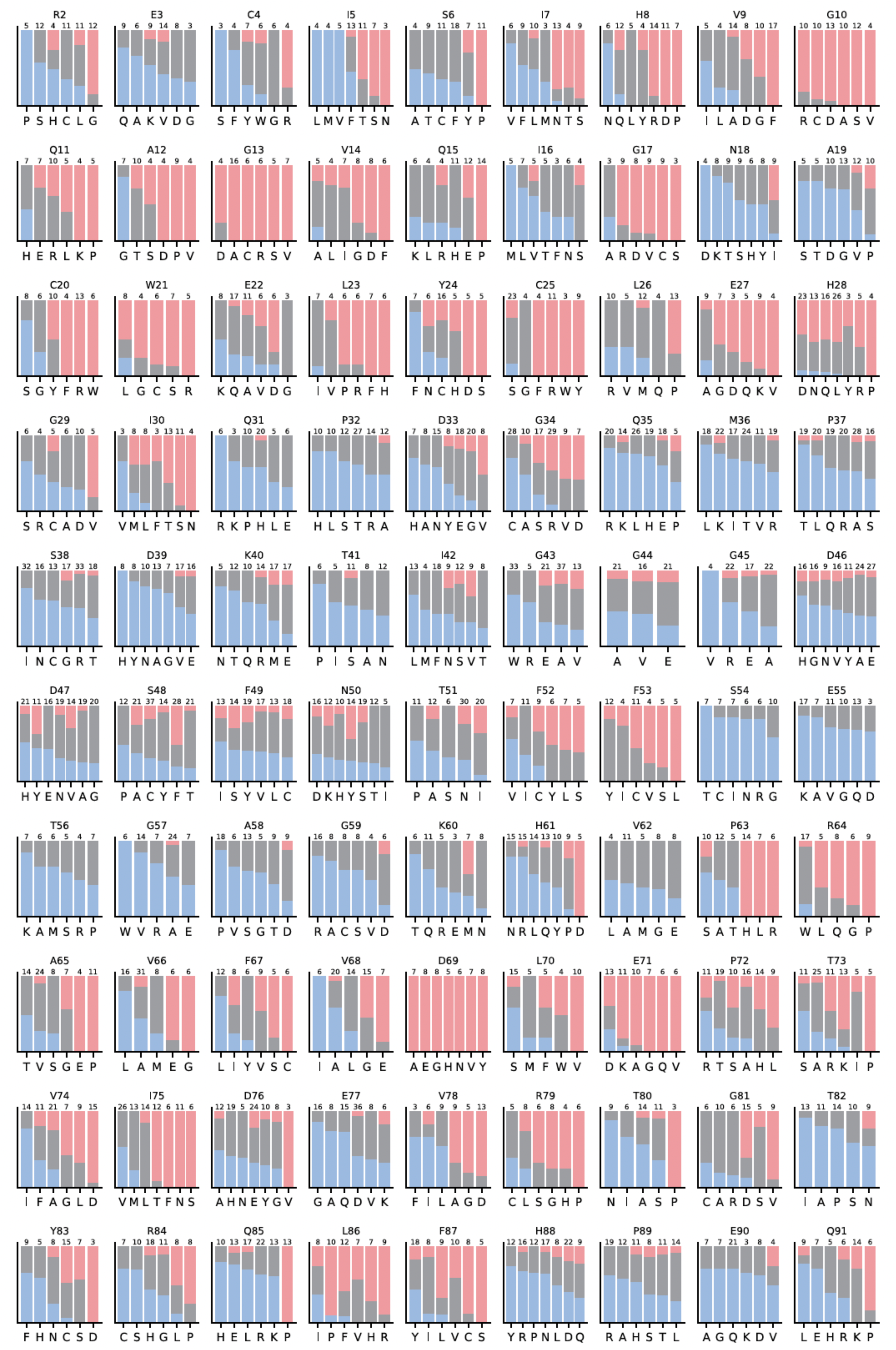
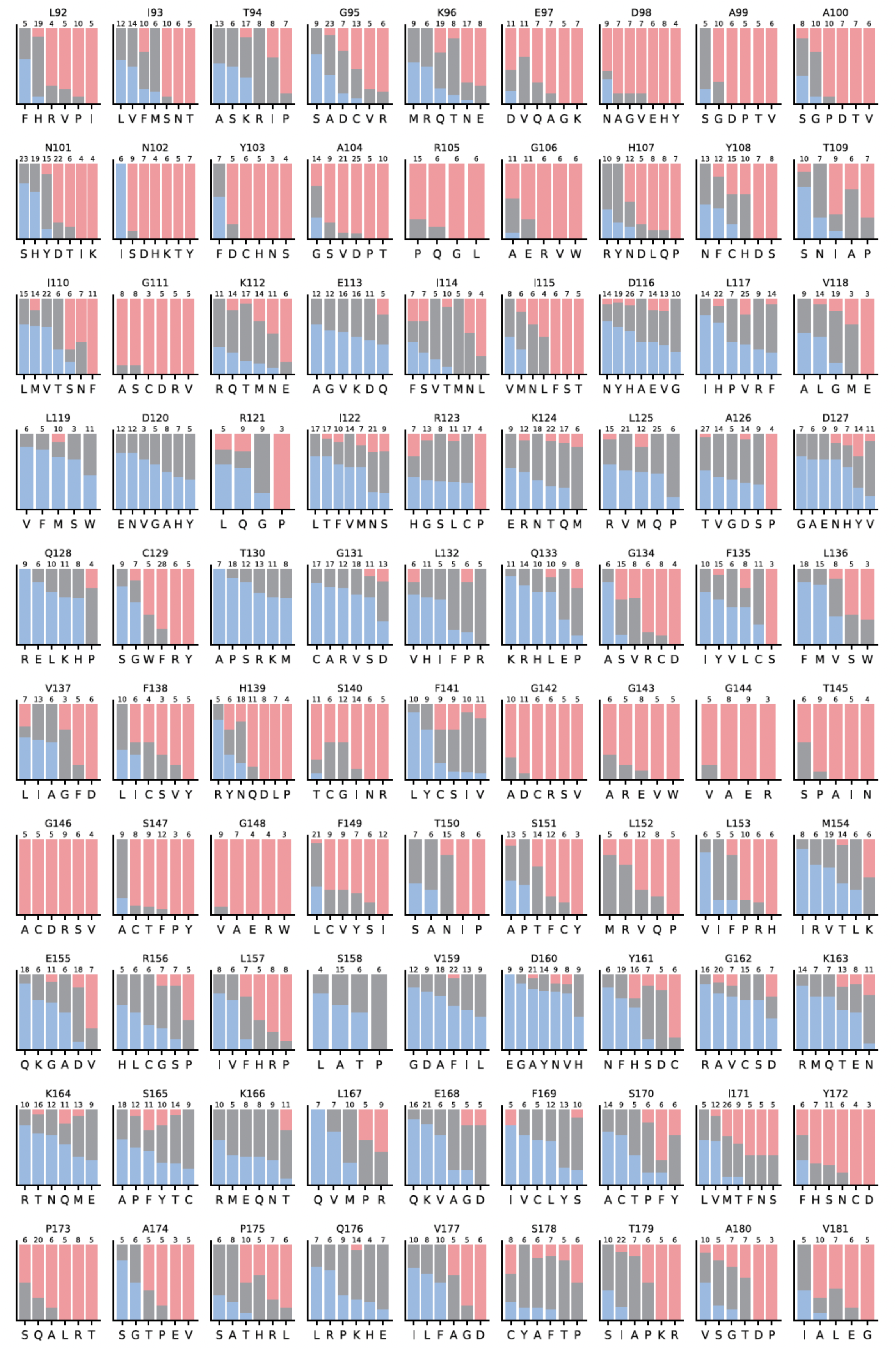
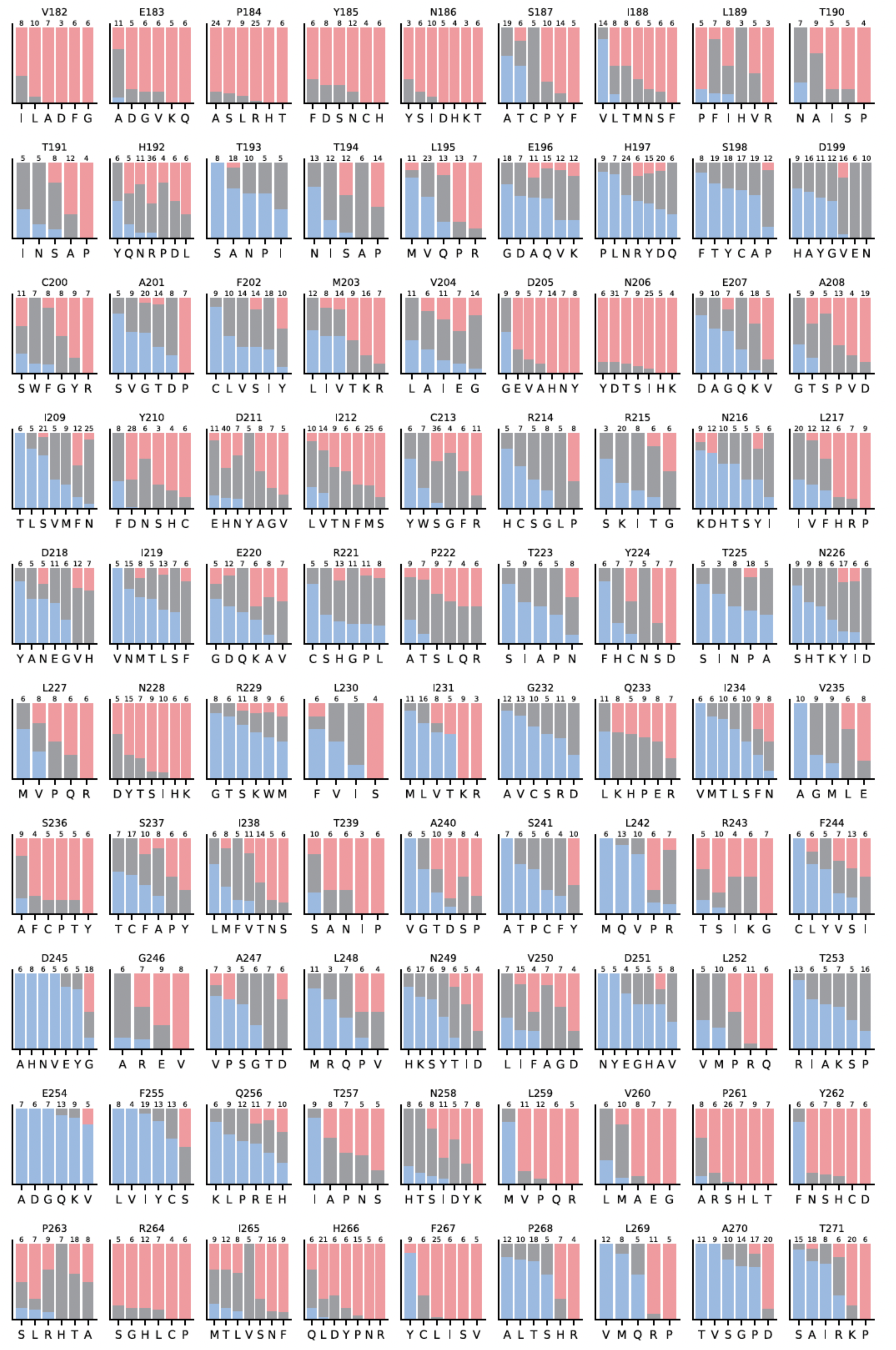
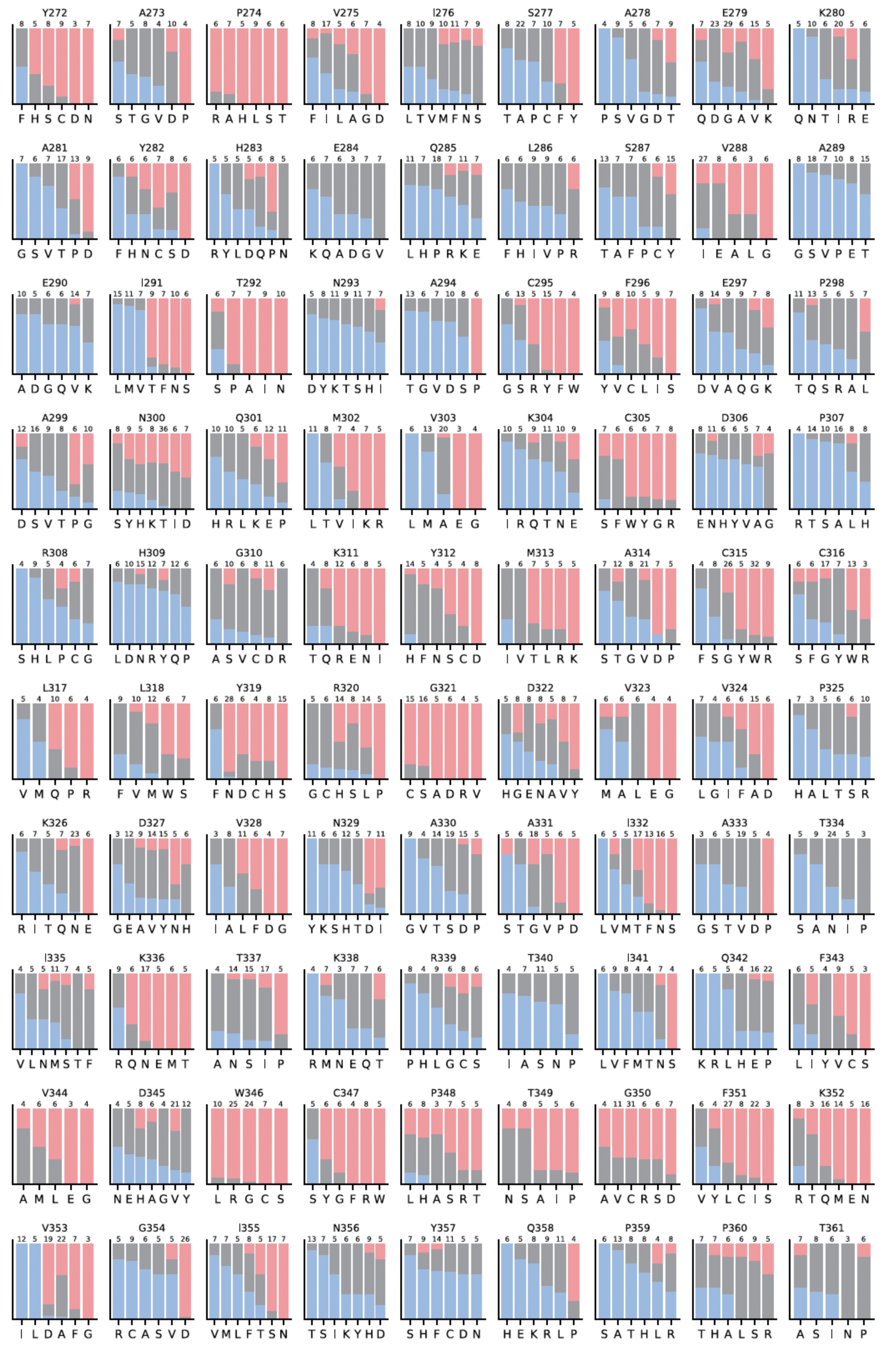
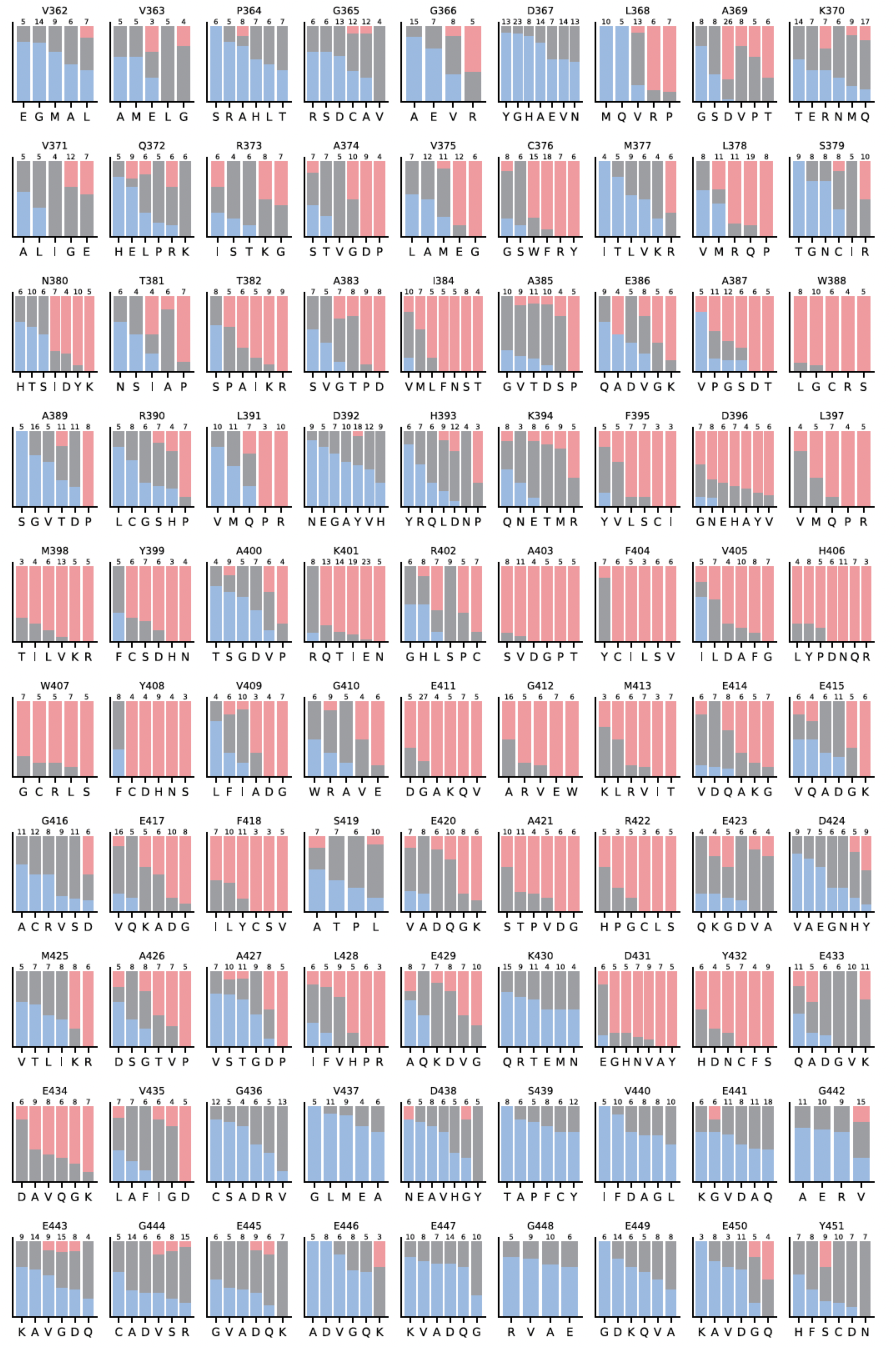
Bargraph representation showing detailed classification results for each TUBA1A SNV, relates to Figure 2. Each panel represents one amino acid residue, which is indicated above the panel. For single nucleotide variants (SNVs), one amino acid could be mutated to 3-7 other forms. Targeted mutations are indicated underneath each panel. Percentage of assembled (A) cells are colored in blue, partially assembled (PA) cells are colored in gray, not assembled (NA) cells are colored in red. The total percentage of cells for each SNV is 100 %. The numbers of cells for assembly score calculation were indicated above the bars of each SNV.

**Figure S3.**
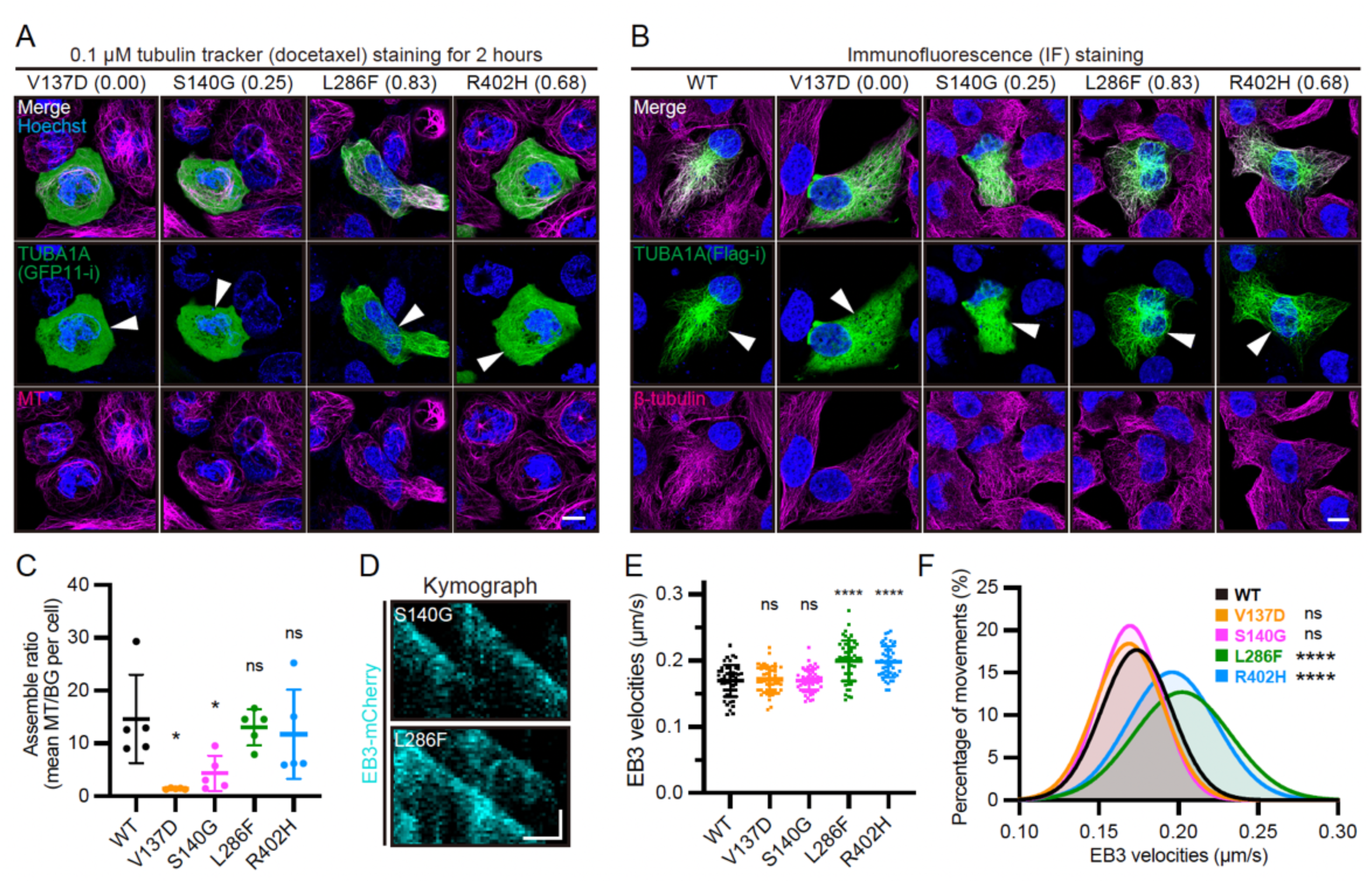
High-content imaging (HCI) results align with immunofluorescence (IF) staining results, relates to Figure 2. (A) Representative images showing localization of split-GFP labeled TUBA1A SNVs in HeLa cells. All variants are associated with tubulinopathies. GFP11-i labeled variants with GFP1-10 were co-transfected into HeLa cells. MTs were stained using 0.1x Tubulin Tracker Deep Red for 2 hours before imaging. All conditions are identical to HCI (Method S1). Nuclei were stained by Hoechst 33342. Transfection-positive cells are indicated by white arrowheads. Scale bar, 10 μm. (B) Representative images showing localization of Flag-tagged TUBA1A SNVs in HeLa cells. Flag tags were inserted into H1-S2 loops of TUBA1A. TUBA1A variants and MTs were visualized by IF staining using Anti-Flag or Anti-β-tubulin first antibodies, individually. Transfection-positive cells are indicated by white arrowheads. Scale bar, 10 μm. (C) TUBA1A (Anti-Flag) fluorescence ratio of MTs to cytosol backgrounds (BG) in each group in (B). Data are shown as mean ± SD. N = 5 cells. For each cell, the average ratio of 10 MTs to their respective background fluorescence is calculated. Unpaired *t* test was performed. (D) Representative kymographs showing EB3-mCherry movements in HeLa cells expressing TUBA1A (S140G) (top) or TUBA1A (L286F) (bottom) variants. Scale bar (horizontal), 2 μm; Scale bar (vertical), 10 sec. (E) EB3-mCherry velocities in HeLa cells expressing each TUBA1A variant. Data are shown as mean ± SD. N = 50 particles. Unpaired *t* test was performed. (F) Distribution of EB3 velocities in each group in (E). Data are shown as Gaussian curves.

**Figure S4.**
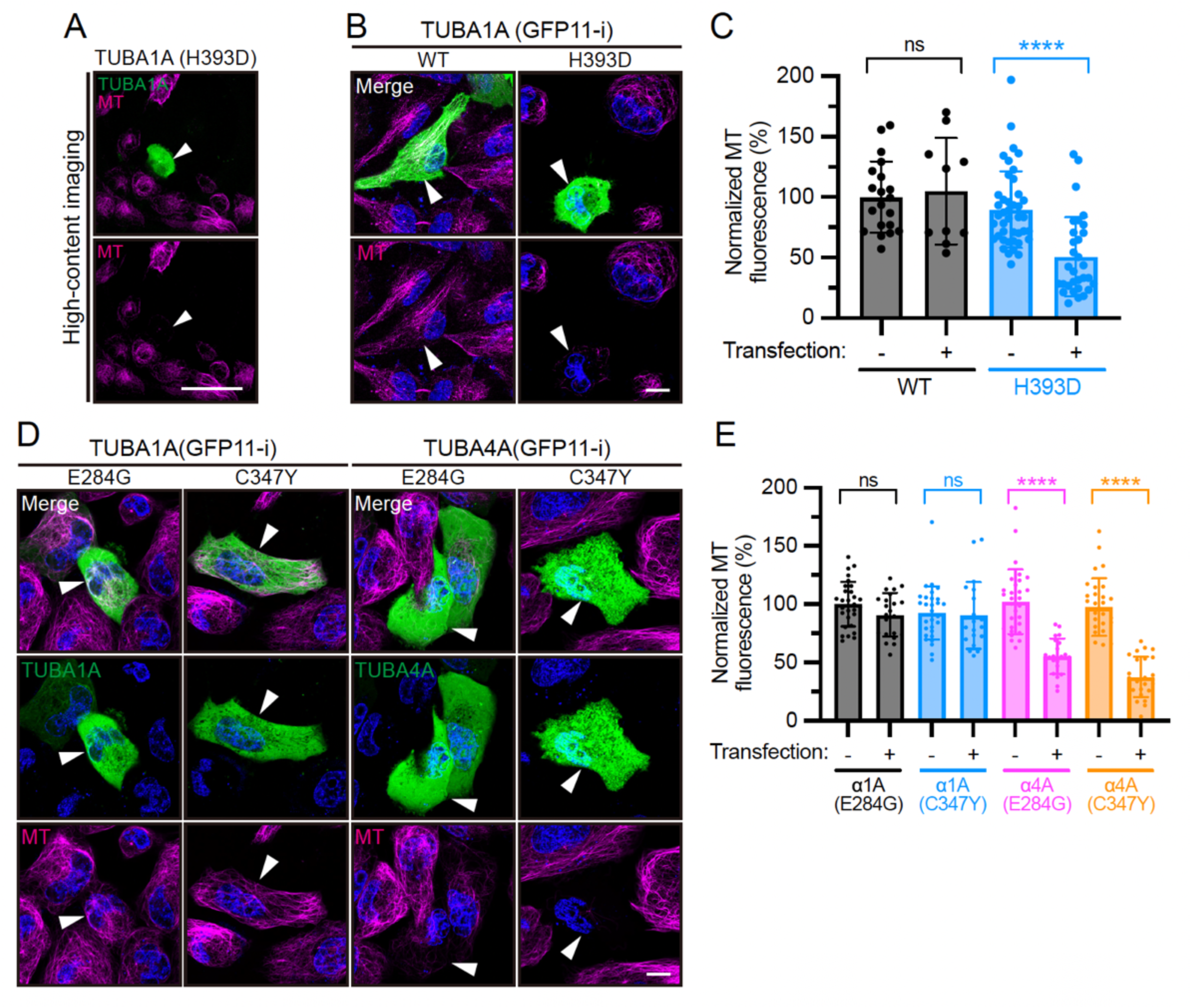
High-content imaging reveals TUBA1A variants that disrupt microtubule architecture in HeLa cells, relates to Figure 2. (A) Representative HCI images of HeLa cells expressing the TUBA1A (H393D) variant. Scale bar, 50 μm. (B) Super-resolution live images of HeLa cells expressing split-GFP labeled wild-type TUBA1A or H393D variant. MTs were stained by 0.1x Tubulin Tracker. Transfection-positive cells are indicated by white arrowheads. Scale bar, 10 μm. (C) Quantification of whole-cell average MT fluorescence intensity for each group in (B). Data are shown as mean ± SD. Mean value of transfection-negative (-) cells in WT group is normalized to 100 %. N > 10 cells per group. (D) Representative images of cells expressing split-GFP labeled TUBA1A (E284G), or TUBA1A (C347Y), or TUBA4A (E284G), or TUBA4A (C347Y) variants. Transfection-positive cells are indicated by white arrowheads. Scale bar, 10 μm. (E) Quantification of whole-cell average MT fluorescence intensity for each group in (D). Mean value of transfection-negative (-) cells in TUBA1A (E284G) group is normalized to 100 %. N > 20 cells per group.

**Figure S5.**
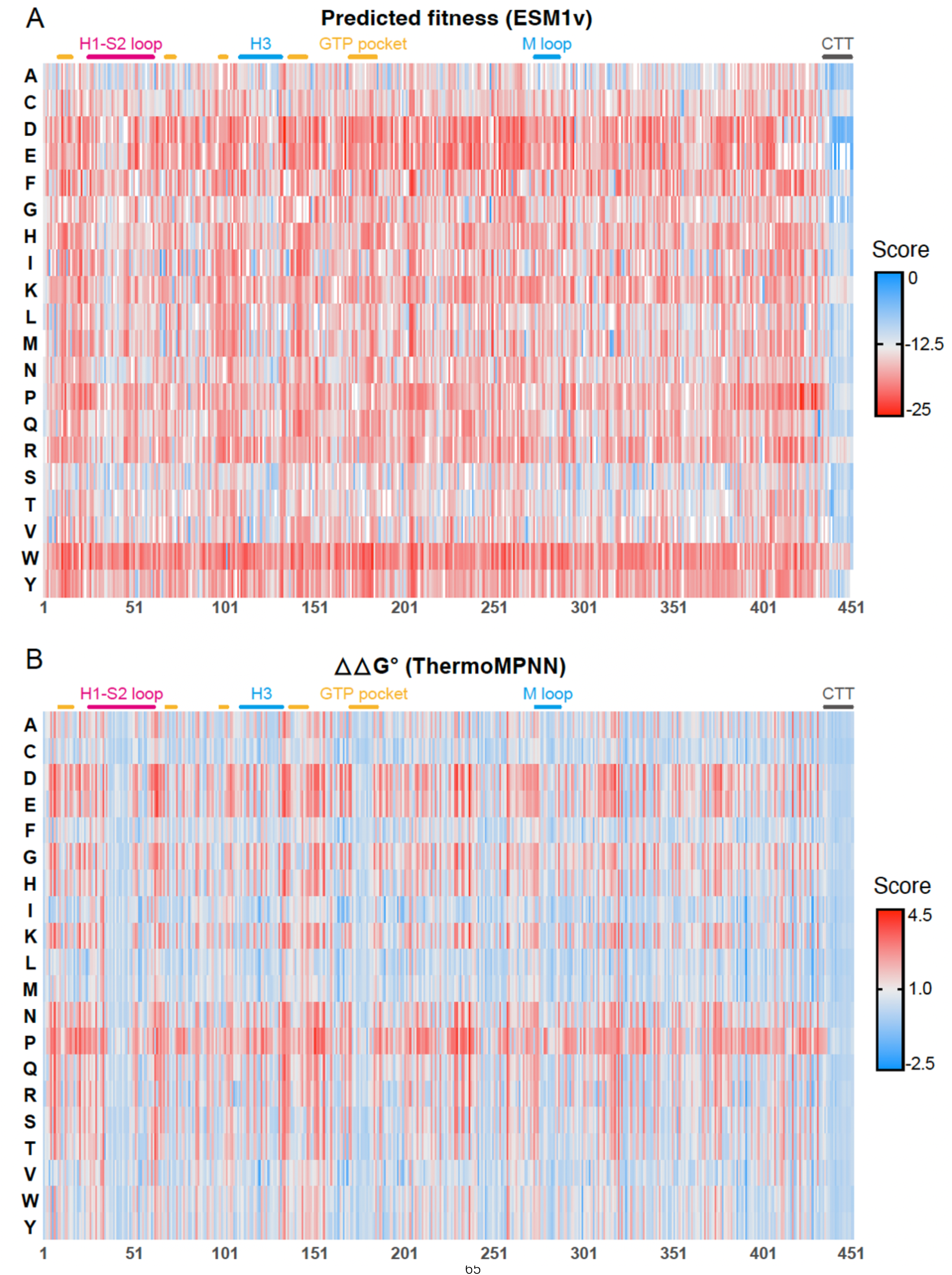
Heatmap representation of predicted fitness (A) by ESM1v, or predicted △△G° (B) by ThermoMPNN for all TUBA1A missense variants, relates to Figure 2. Regions corresponding to H1-S2 loop, or H3 helix, or GTP-binding pocket, or M loop, or C-terminal tail (CTT) are differentially colored and indicated above each heatmap.

**Figure S6.**
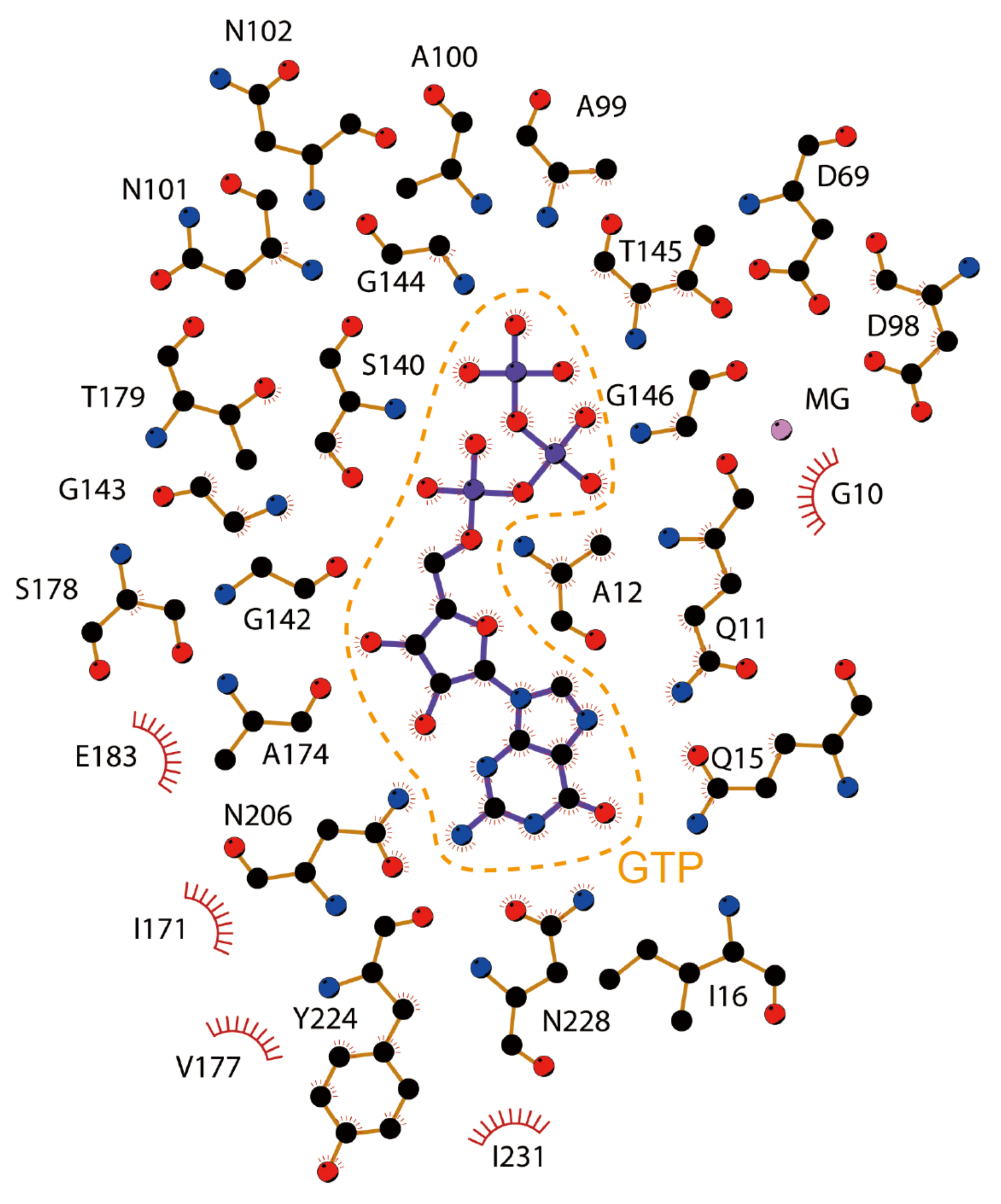
Diagram showing interaction between GTP and neighboring amino acid residues in TUBA1A, relates to Figure 3. GTP is indicated by orange dashed lines. Diagram is produced using LigPlot^+^ v.2.2 (http://www.ebi.ac.uk/thornton-srv/software/LigPlus/).

**Figure S7.**
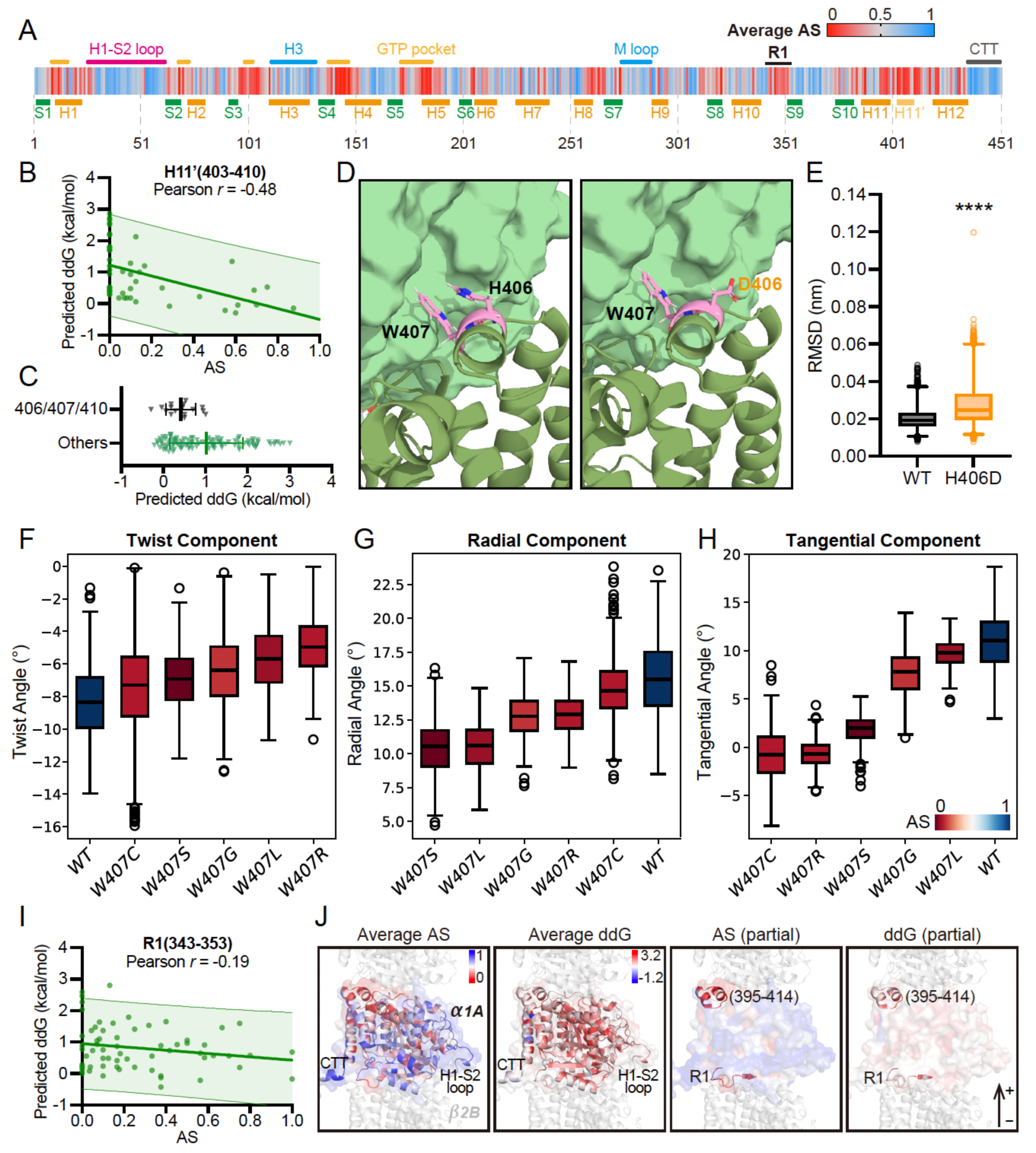
Assembly score landscape and molecular dynamics simulation of TUBA1A reveals critical regions mediating intradimer and interdimer interactions, relates to Figure 4. (A) Heatmap representation showing average AS of TUBA1A residues. Functional domains (H1-S2 loop, H3, GTP pocket, M loop or C-terminal tail) are indicated above the heatmap, and region 1 (R1) is indicated by black lines. Helixes and β-strands are indicated below the heatmap. (B) Correlation scatter plot of AS and predicted △△G° for all SNVs in helix 11’ (H11’, residue 402-410). 95 % prediction band was shown. (C) Predicted △△G° distribution of SNVs in H406/W407/G410, or other residues in H11’. Data are shown as mean ± SD. (D) Representative intradimer face of wild-type heterodimer (left) or H406D mutant (right). TUBA1A was shown as cartoon and colored in dark green. TUBB2B was shown as surface and colored in light green. (E) Quantitative statistics of the root mean square deviation (RMSD) of H406 residue (WT) or D406 residue (H406D) during simulation. Sampling is conducted every 2 ns in the last 500 ns of the 1-μs simulation. (F-H) Quantitative statistics of twist angles (F), radial bending angles (G), or tangential bending angles (H) of wild-type or W407 variants during simulation. A single tubulin heterodimer is employed for MDS (see Methods). Sampling is conducted every 2 ns in the last 500 ns of the 1-μs simulation. (I) Correlation scatter plot of AS and predicted △△G° for all SNVs in R1 (residues 343-353). (J) 3D structural representation of average AS and average predicted △△G° in TUBA1A. In the two rightmost panels, R1 and residues 395-414 are highlighted in TUBA1A structure. +, MT plus end.

**Figure S8.**
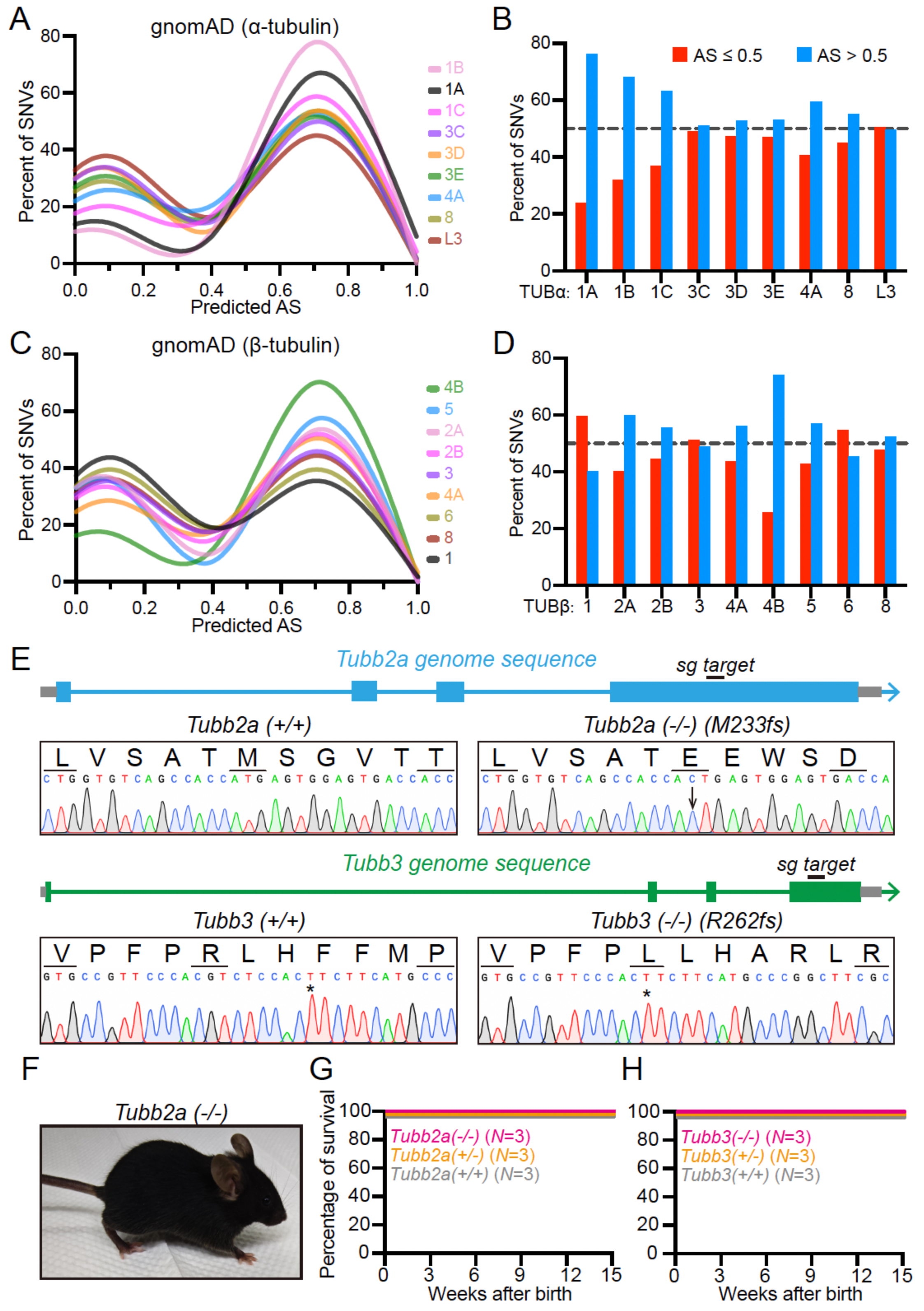
Loss-of-function landscape of TUBA1A highlights differential constraints across tubulin isotypes, relates to Figure 7. (A) Frequency distribution of predicted AS in each α-tubulin isotype in Figure 7A. The total percentage of each group is 100 %. (B) Bargraph representation showing relative proportion of SNVs with AS ≤ 0.5 (red) or AS > 0.5 (blue) in each α-tubulin isotype in Figure 7A. (C) Frequency distribution of predicted AS in each β-tubulin isotype in Figure 7B. (D) Bargraph representation showing relative proportion of SNVs with AS ≤ 0.5 (red) or AS > 0.5 (blue) in each β-tubulin isotype in Figure 7B. (E) Peak diagrams showing Sanger genotyping results of *Tubb2a* (+/+) and *Tubb2a* (-/-) mice (top), or *Tubb3* (+/+) and *Tubb3* (-/-) mice (bottom). Frameshift mutation occurs in *Tubb2a* (-/-) mice through insertion of one nucleotide (C, indicated by a black arrow). Frameshift mutation occurs in *Tubb3* (-/-) mice through deletion of eight nucleotides (GTCTCCAC), and the following nucleotide (T) is indicated by a black asterisk. Diagrams of the *Tubb2a* and *Tubb3* genomic loci are shown above the Sanger genotyping results, with sgRNA target sites highlighted. (F) Representative image of an 8-week-old female C57BL/6 mouse of *Tubb2a* (-/-) genotype. (G and H) Kaplan–Meier survival curves showing *Tubb2a* (G) or *Tubb3* (H) knock-out mice are viable.

**Movie S1. Molecular dynamics simulation depicting the interaction between GTP and TUBA1A (WT) or A12D mutant.** This movie corresponds to Figure 3F. Total simulation time: 1,00 ns.

**Movie S2. Molecular dynamics simulation illustrating protofilament dynamics of TUBA1A (WT), or M398I variant, or F351I variant, or L397R variant.** This movie corresponds to Figure 4D–4G. Total simulation time: 1,000 ns.

