## Supplementary material for "A comprehensive functional landscape of α-tubulin TUBA1A variants illuminates microtubule biology and refines clinical classification": Method S1

**Method S1: Detailed protocols and parameters for implementation of the integrated experimental-computational platform for comprehensive mutagenesis**

**Content:**

**1. The *tuba1a (gfp11-i)* coding-domain sequence and primer sequences (Page 2-3)**

**2. Algorithm for automatic primer design (Page 4)**

**3. Protocol for modified SOEing-PCR (Page 5-8)**

**4. Protocol for transfection of SOEing-PCR products (Page 9)**

**5. Protocol for high-content imaging (HCI) (Page 10)**

**6. Manual proofreading of assembly score (AS) (Page 11)**

**1. The *tuba1a (gfp11-i)* coding-domain sequence and primer sequences**

Sequence of CMV enhancer – CMV promoter – *tuba1a (gfp11-i)* – bGH terminator:

GACATTGATTATTGACTAGTTATTAATAGTAATCAATTACGGGGTCATTAGTTCATAGCCCATATATGGAGTTCCGCGTTACATAACTTACGGTAAATGGCCCGCCTGGCTGACCGCCCAACGACCCCCGCCCATTGACGTCAATAATGACGTATGTTCCCATAGTAACGCCAATAGGGACTTTCCATTGACGTCAATGGGTGGACTATTTACGGTAAACTGCCCACTTGGCAGTACATCAAGTGTATCATATGCCAAGTACGCCCCCTATTGACGTCAATGACGGTAAATGGCCCGCCTGGCATTATGCCCAGTACATGACCTTATGGGACTTTCCTACTTGGCAGTACATCTACGTATTAGTCATCGCTATTACCATGGTGATGCGGTTTTGGCAGTACATCAATGGGCGTGGATAGCGGTTTGACTCACGGGGATTTCCAAGTCTCCACCCCATTGACGTCAATGGGAGTTTGTTTTGGCACCAAAATCAACGGGACTTTCCAAAATGTCGTAACAACTCCGCCCCATTGACGCAAATGGGCGGTAGGCGTGTACGGTGGGAGGTCTATATAAGCAGAGCTCTCTGGCTAACTAGAGAACCCACTGCTTACTGGCTTATCGAAATTAATACGACTCACTATAGGGAGACCCAAGCTTGGTACCGAGCTCGGATCCGCCACC**ATG**CGTGAGTGCATCTCCATCCACGTTGGCCAGGCTGGTGTCCAGATTGGCAATGCCTGCTGGGAGCTCTACTGCCTGGAACACGGCATCCAGCCCGATGGCCAGATGCCAAGTGACAAGACCATTGGGGCTGGAAGCGGGAGCGGAGCCGGTAGCGGTAGCGGCGCAGGGAGCGGGCGAGATCATATGGTACTTCATGAATATGTAAATGCAGCTGGTATAACTGCTGGAAGTGGTAGCGGGGCTGGAAGTGGTAGCGGAGCGGGCAGTGGAGGAGGAGATGATTCCTTCAACACCTTCTTCAGTGAGACGGGGGCTGGCAAGCATGTGCCCCGGGCAGTGTTTGTAGACTTGGAACCCACAGTCATTGATGAAGTTCGCACTGGCACCTACCGCCAGCTCTTCCACCCTGAGCAACTTATCACAGGCAAGGAAGATGCTGCCAATAACTATGCCCGAGGGCACTACACCATTGGCAAGGAGATCATTGACCTCGTGTTGGACCGAATTCGCAAGCTGGCCGACCAGTGCACGGGTCTCCAGGGCTTCTTGGTTTTCCACAGCTTTGGTGGGGGAACTGGTTCTGGGTTCACCTCCCTGCTCATGGAACGTCTCTCAGTTGATTATGGCAAGAAGTCCAAGCTGGAGTTCTCTATTTACCCGGCGCCCCAGGTTTCCACAGCTGTAGTTGAGCCCTACAACTCCATCCTCACCACCCACACCACCCTGGAGCACTCTGATTGTGCCTTCATGGTAGACAATGAGGCCATCTATGACATCTGTCGTAGAAACCTCGATATTGAGCGTCCAACCTATACTAACCTGAATAGGTTAATAGGTCAAATTGTGTCCTCCATCACTGCTTCCCTGAGATTTGATGGAGCCCTGAATGTTGACCTGACAGAATTCCAGACCAACCTGGTGCCCTATCCCCGCATCCACTTCCCTCTGGCCACATATGCCCCTGTCATCTCTGCTGAGAAAGCCTACCATGAACAGCTTTCTGTAGCAGAGATCACCAATGCTTGCTTTGAGCCAGCCAACCAGATGGTGAAATGTGACCCTCGCCATGGTAAATACATGGCTTGCTGCCTGTTGTACCGTGGTGACGTGGTTCCCAAAGATGTCAATGCTGCCATTGCCACCATCAAGACCAAGCGTACCATCCAGTTTGTGGATTGGTGCCCCACTGGCTTCAAGGTTGGCATCAACTACCAGCCTCCCACTGTGGTGCCTGGTGGAGACCTGGCCAAGGTACAGAGAGCTGTGTGCATGCTGAGCAACACCACAGCCATTGCTGAGGCCTGGGCTCGCCTGGACCACAAGTTTGACCTGATGTATGCCAAACGTGCCTTTGTTCACTGGTACGTTGGGGAGGGGATGGAGGAAGGTGAGTTTTCAGAGGCCCGTGAGGACATGGCTGCCCTTGAGAAGGATTATGAGGAGGTTGGTGTGGATTCTGTTGAAGGAGAGGGTGAGGAAGAAGGAGAGGAATAC**TAA**TCTAGAGGGCCCTATTCTATAGTGTCACCTAAATGCTAGAGCTCGCTGATCAGCCTCGACTGTGCCTTCTAGTTGCCAGCCATCTGTTGTTTGCCCCTCCCCCGTGCCTTCCTTGACCCTGGAAGGTGCCACTCCCACTGTCCTTTCCTAATAAAATGAGGAAATTGCATCGCATTGTCTGAGTAGGTGTCATTCTATTCTGGGGGGTGGGGTGGGGCAGGACAGCAAGGGGGAGGATTGGGAAGACAATAGCAGGCATGCTGGGGATGCGGTGGGCTCTATGG

Sequence of the upstream general primer pF1 (as shown in Figure 1C), with pF2 sequence in blue and template-binding sequence underlined:

GGCCACGCTACCATGGAGCTCCAAATAATGCGCGTTGACATTGATTATTGACTAG

Sequence of the upstream general primer pF2 (as shown in Figure 1C):

CCATGGAGCTCCAAATAATG

Sequence of the downstream general primer pR1 (as shown in Figure 1C), with pR2 sequence in purple and template-binding sequence underlined:

CGATGAGCATGATTTGACGTCATGAGAGGCCCATAGAGCCCACCGCATCCC

Sequence of the downstream general primer pR2 (as shown in Figure 1C):

GATTTGACGTCATGAGAGGC

**2. Algorithm for automatic primer design**

**Input:** mutated position P, which will be mutated to M, original sequence S

**Output:** forward and reverse primers F and R.

1. S[P] = M
2. # define seed region and split seed into frag1 and frag2.

seed = S[P-25:P+25]

frag1, frag2 = seed[:25], seed[26:]

\# Tm temperature optimization on frag1.

frag1_index = 25

while Tm(frag1) < 55℃:

seed expand 1 bp at 3’;

frag1_index += 1;

frag1 = seed[:frag1_index];

while Tm(frag) > 60℃:

trim the seed 1 at 3’;
 frag1_index -= 1;

frag1 = seed[:frag1_index];

if len(frag1) <= 20:

break

3. #Ensure the 3’ terminal of frag1 is ‘G/C’ when the Tm is within the range

frag1_c = frag1[first_c(frag1):]

frag1_g = frag1[first_g(frag1):]

frag1_candidate = [n for n in [frag1, frag1_c, frag1_g] if Tm(n) $\in(55,60)$]

frag1 = ${Argmax}_{Tm}$(frag1_candidate)

4. #Reverse complicate to create F and R

R = seed

F = ReverseComplicate(seed)

5. #Trim frag2 to make sure that there are around 24 bp overlap

Trim_bases = int(len(F)-24)/2

R = R[Trim_bases:]

F = F[Trim_bases:]

return F, R

**3. Protocol for modified SOEing-PCR**

As shown in Figure 1C, circular plasmids carrying the enhancer / promoter, coding-domain sequence (CDS) of gene of interest (GOI) such as human α-tubulin *tuba1a*, and the terminator are employed as templates. Fluorescent tags (such as split-sfGFP or EGFP) or affinity tags (such as HA or Flag tag) could be fused to POI directly, as this will not affect the efficiency of modified SOEing-PCR.

For each mutation to be constructed, a pair of forward and reverse primers (namely pMut-F and pMut-R) were designed (Figure 1B). The pMut-F binds to the complementary chain of the template plasmid, and therefore is the counterpart of the general primer pR1 (which binds to the downstream site of terminator). As a result, the PCR fragment corresponding to the latter part of GOI is amplified using the pMut-F / pR1 primer pair. The targeted mutation and the pR2-specific binding region are introduced into the resulted PCR fragment, located on the two opposite ends of the fragment. Likewise, the pMut-R binds to the sense chain of the template plasmid, and therefore is the counterpart of the general primer pF1 (which binds to the upstream site of enhancer / promoter). As a result, the PCR fragment corresponding to the former part of GOI is amplified using the pF1 / pMut-R primer pair. The targeted mutation and the pF2-specific binding region are introduced into the resulted PCR fragment, located on the two opposite ends of the fragment.

For each pair of pMut-F / pMut-R, the corresponding amplified latter PCR fragment and former PCR fragment served as the templates together for final amplification of full-length linear PCR products containing targeted mutant. Consequently, to enable high-throughput construction of mutants, primers were typically synthesized in 96-well or 384-well standard PCR plates. Notably, forward primers (or pMut-F) should be synthesized together in a patch of PCR plates in designated orders (for example, well A01 represented the first mutation, well A02 represented the second mutation, well B01 represented the 13^th^ mutation, and well H12 represented the 96^th^ mutation). Reverse primers (or pMut-R) should be synthesized together in another patch of PCR plates in the same order as their corresponding pMut-F (that is, well A01 represented the first mutation, well A02 represented the second mutation…) (Figure 1A). Therefore, to construct all 2,683 *tuba1a* missense SNVs, 28 96-well PCR plates containing 2,683 different pMut-F primers, and 28 96-well PCR plates containing 2,683 different pMut-R primers, were synthesized as the primer pool for subsequent PCR amplification. This large-scale primer synthesis could be directly finished by commercial companies (e.g., Beijing Genomics Institute, or BGI) at the price of 0.03–0.1 US dollar per nucleotide.

For the first two rounds of PCR (namely PCR1 and PCR2) (Figure 1C), the following concentration for each PCR component was employed:

**The 10 μL PCR reaction mix:**

1 x Phanta Max Master Mix (Vazyme, #P525-01)

0.2 ng/μL Plasmid template

0.1 pM General primer pF1 (or pR1)

0.25 pM Mutation-specific primer pMut-R (or pMut-F)

To enable high-throughput preparation of PCR reaction mixtures, the Biomek® Nxp automatic working station (Beckman) was employed for allocation of the PCR components above. As all mutation-specific primers (pMut-R or pMut-F) has already been synthesized in 96-well PCR plates, this process could be simply finished by transferring 2.5 μL of 1 pM pMut-R (or pMut-F) to 7.5 μL of pre-distributed PCR reaction mix containing enzyme, substrates, templates and the general primer pF1 (or pR1). The following PCR condition was employed:

| Step | Temperature | Time | Cycle |
| --- | --- | --- | --- |
| Denaturing | 95 ℃ | 3 min | 1 |
| Denaturing | 95 ℃ | 15 sec | 25 |
| Annealing | 58 ℃ | 15 sec |  |
| Extension | 72 ℃ | 30 sec – 90 sec |  |
| Extension | 72 ℃ | 5 min | 1 |

For the last round of PCR (namely PCR3) (Figure 1C), the following concentration for each PCR component was employed:

**The 30 μL PCR reaction mix:**

1 x Phanta Max Master Mix (Vazyme, #P525-01)

Linear PCR product from PCR1 (1:400 diluted in final reactions)

Linear PCR product from PCR2 (1:400 diluted in final reactions)

0.4-1 pM General primer pF2

0.4-1 pM General primer pR2

The Biomek® Nxp automatic working station (Beckman) was employed for dilution of linear PCR products from PCR1 or PCR2, and subsequent transferring of them into pre-distributed PCR reaction mix containing enzyme, substrates and the general primer pF2 and pR2. The following condition for PCR amplification was employed:

| Step | Temperature | Time | Cycle |
| --- | --- | --- | --- |
| Denaturing | 95 ℃ | 3 min | 1 |
| Denaturing | 95 ℃ | 15 sec | 33 |
| Annealing | 53 ℃ | 15 sec |  |
| Extension | 72 ℃ | 2 min |  |
| Extension | 72 ℃ | 5 min | 1 |

For construction of all 2,683 *tuba1a* SNVs, the modified SOEing-PCR (as shown in Figure 1C) could be accomplished within 1-2 days. Agarose gel electrophoresis could help to ensure the successful and specific amplification of linear PCR products. Sanger sequencing could help to ensure the successful and specific mutagenesis. The SOEing-PCR products could be used directly for transfection or stored at -20 ℃ for long-term conservation.

**4. Protocol for transfection of SOEing-PCR products**

Typically, the concentrations of linear DNA products in final reaction mixes would be approximately 100 ng/μL per sample. DNA purification is not required before transfection, as we found this process does not help to improve transfection efficiency. For large-scale transfection, polyethylenimine (PEI) MW40,000 transfection reagent (Yeasen) was employed. The following protocol could be utilized for transfection:

1. Dilute 1.5 μL SOEing-PCR product and 150 ng pCDNA3.0-*gfp1-10* plasmid in 5 μL serum-free opti-MEM (Gibco) per sample
2. Dilute 0.6 μL PEI transfection reagent (1 mg/mL) in 5 μL opti-MEM per sample
3. Mix the PEI solution (step 2) and the DNA solution (step 1) together per sample by pipetting gently and incubate for 15 min at room temperature
4. Directly add the transfection mixtures (step 3) per sample into each well of cells seeded in 96-well cell culture plates with appropriate density
5. Incubate the cells at 37 ℃ with 5 % CO_2_ for 12 – 24 hours before live-cell imaging

**5. Protocol for high-content imaging (HCI)**

Images were acquired using the ImageXpress® Confocal HT.ai system (Molecular Devices) with a 60×/1.4 objective, a 37 ℃ and 5 % CO_2_ chamber, a 60-μm pinhole camera, the autofocus mode and Cy5 / FITC channels. We selected the best z-plane which displays the clearest MT networks in Cy5 channel (the channel for Tubulin Tracker^TM^ Deep Red) as the reference plane for both Cy5 and FITC channels. The AI mode of this confocal system enabled autofocus at every imaging site to find the designated reference planes, which reflected the clearest MT networks. As a result, all images from every site were automatically taken at the best resolution, making it feasible for subsequent deep learning-based MT recognization. To stain MTs, the cultural medium was replaced with 40 μL DMEM containing 0.1x Tubulin Tracker^TM^ Deep Red for each well (in 96-well plates). High-content imaging was performed after incubating the cells at 37 ℃ with 5 % CO_2_ for 30 min. The laser intensities were set to 100 %, and exposure time was 200 ms for Cy5 channel, and 500 ms for FITC channel (GFP channel). 36 different sites were chosen for unbiased imaging per well (each well represents one SNV), which enables acquiring enough transfection-positive cells for subsequent analysis. Typically, the imaging process for one 96-well plate could be finished within 1.5-2 hours, which does not impact the cellular phenotype of GFP11-i labeled tubulins or MTs significantly (Figure S3A). In total, 193,176 raw images were acquired (2683 wells × 36 sites per well × 2 channels).

**6. Manual proofreading of assembly score (AS)**

After automatic classification of all transfection-positive cells, proofreading was performed by examining all cells with labels. Label of one cell could be manually changed to another if another classification is preferred in visualization. After proofreading, SNVs with less than 3 authorized cells (those with an ‘Assemble’ or ‘Partially Assemble’ or ‘Not Assemble’ label) were screened out to perform HCI again. After the second round of HCI scanning, only 3 (of 2683) SNVs had less than 3 authorized cells. Through agarose gel electrophoresis analysis, we concluded that the modified SOEing-PCR products of these 3 SNVs did not contain targeted full-length DNA products. We solved this problem by manually constructing plasmids for site-directed mutagenesis of the 3 mutants, as the traditional methodology did. Consequently, we achieved 99.9 % (2680 / 2683) success rate in modified SOEing-PCR process.

The assembly score (AS) was subsequently quantified by a weighted average calculation based on the classification results:

$$AS=\frac{A+0.5PA}{A+PA+NA}$$
